# Genetic control of pluripotency epigenome determines differentiation bias in mouse embryonic stem cells

**DOI:** 10.1101/2021.01.15.426861

**Authors:** Candice Byers, Catrina Spruce, Haley J. Fortin, Anne Czechanski, Steven C. Munger, Laura G. Reinholdt, Daniel A. Skelly, Christopher L. Baker

## Abstract

Genetically diverse pluripotent stem cells (PSCs) display varied, heritable responses to differentiation cues in the culture environment. By harnessing these disparities through derivation of embryonic stem cells (ESCs) from the BXD mouse genetic reference panel, along with C57BL/6J (B6) and DBA/2J (D2) parental strains, we demonstrate genetically determined biases in lineage commitment and identify major regulators of the pluripotency epigenome. Upon transition to formative pluripotency using epiblast-like cells (EpiLCs), B6 quickly dissolves naïve networks adopting gene expression modules indicative of neuroectoderm lineages; whereas D2 retains aspects of naïve pluripotency with little bias in differentiation. Genetic mapping identifies 6 major *trans*-acting loci co-regulating chromatin accessibility and gene expression in ESCs and EpiLCs, indicating a common regulatory system impacting cell state transition. These loci distally modulate occupancy of pluripotency factors, including TRIM28, P300, and POU5F1, at hundreds of regulatory elements. One *trans*-acting locus on Chr 12 primarily impacts chromatin accessibility in ESCs; while in EpiLCs the same locus subsequently influences gene expression, suggesting early chromatin priming. Consequently, the distal gene targets of this locus are enriched for neurogenesis genes and were more highly expressed when cells carried B6 haplotypes at this Chr 12 locus, supporting genetic regulation of biases in cell fate. Spontaneous formation of embryoid bodies validated this with B6 showing a propensity towards neuroectoderm differentiation and D2 towards definitive endoderm, confirming the fundamental importance of genetic variation influencing cell fate decisions.

## Introduction

The ability to form all somatic and germline tissues, while maintaining the capacity to self-renew, are defining features of PSCs^1–4^. Harnessing this potential will transform regenerative medicine. Yet, most studies seeking to identify mechanisms that underlie acquisition of pluripotency and fate determination utilize cells with limited genetic diversity. Historically, successful derivation of naïve mouse ESCs was achieved for permissive strains (i.e. 129 and C57BL/6 lineages). Nonpermissive strains, such as D2, required inhibition of differentiation pathways (i.e. 2i)^5–7^, demonstrating that the establishment and maintenance of pluripotency are intrinsic to genetic background. Further, success in derivation and differentiation of human induced PSCs (hiPSCs) and ESCs (hESCs) are influenced by genetic background^8–13^. While we^14^ and others^15^ have recently investigated how genetic variation impacts pluripotency, less is understood about how genetic backgrounds influence cell state transitions.

During development, cells within the epiblast progress along a pluripotent spectrum from pre- to post-implantation^16,17^, which can be modeled *in vitro*^18^. ESCs derived from the preimplantation epiblast (E4.5) capture naïve pluripotency^1,2,19^, while epiblast stem cells (EpiSCs) derived post-implantation (E6.5) represent primed pluripotency^20^. Between these two states exists formative pluripotency. Transit through formative pluripotency is required for multi-lineage competency^21–23^, and this state can be modeled *in vitro* by EpiLCs^21^. While culture conditions can provide external signaling cues that positions a cell along the pluripotency spectrum^22^, variability in acquisition of desired pluripotent states is, in part, dependent on the ESC’s strain of origin^6,14,15,24–27^. Importantly, differentiation propensity to specific germ layers has been linked to the origin of ESCs within this pluripotency spectrum^28^. This demonstrates interactions between genetic background and media conditions may drive cell fate.

Reconfiguration of chromatin accompanies, and often precedes^29,30^, transcriptomic changes during cell state transitions^31–35^. Regulatory elements, such as enhancers and promoters, largely act in *cis* to locally control developmentally programmed gene networks^36^. In concert, DNA binding proteins act in *trans* at many *cis*-regulatory elements to either activate or repress gene expression. During development, dramatic alteration of the regulatory landscape can occur even in the absence of changes in transcription factor (TF) expression. For example, when naïve ESCs transition to formative EpiLCs, expression of *Pou5f1/Oct4* remains high, while occupancy of POU5F1 at regulatory elements is globally rewired^37,38^, suggesting the existence of a set of unknown chromatin regulators that influence cell state transitions. Here, we combine the strength of systems genetics, utilizing extant variation in a diverse genetic reference population, with the power of modeling developmental transitions using PSCs, to identify loci that govern chromatin and gene regulation during exit from pluripotency and determine differentiation propensity.

## Results

### Position along the pluripotency spectrum is determined by genetic background

To assess genetic control over pluripotency and differentiation, we derived three biological replicate ESCs from independent blastocysts using B6 and D2 mice. Success in derivation of germline-competent ESCs for these strains required conditions that included serum, feeders, and 2i^6^. Both B6 and D2 ESCs were differentiated to EpiLCs (**Fig. 1a**) and transcript abundance was measured by RNA-sequencing (-seq) and chromatin accessibility by ATAC-seq. While both strains effectively transition from naïve to formative pluripotency, as indicated by expression and accessibility of binding sites of naïve and primed markers, distinct morphological differences were observed (**Fig. 1a-c and Extended Data Fig. 1a&b**). To determine major features of variation, we performed principal-component analysis (PCA) on transcript abundance. While biological replicates closely clustered, demonstrating high reproducibility, PC1 separated ESCs from EpiLCs and PC2 captured strain variation within a cell state (**Fig. 1b**). For ESCs, naïve pluripotency factors (*Klf4, Sox2, Esrrb, Nanog, Pou5f1,* and *Tfcp2l1*) were highly expressed but not differentially abundant between B6 and D2 (**Fig. 1d**). Overall, 1,785 differentially expressed genes (DEGs) in ESCs, and 2,209 DEGs in EpiLCs (FDR < 0.05 and log_2_FC > 1) were identified between B6 and D2 (**Fig. 1e and Extended Data Fig. 1c, Extended Data Table 1**). Of these, only 792 (19.8%) DEGs are shared between cell states, indicating that the strain-dependent DEGs are largely unique to each cell state.

**Figure 1:**
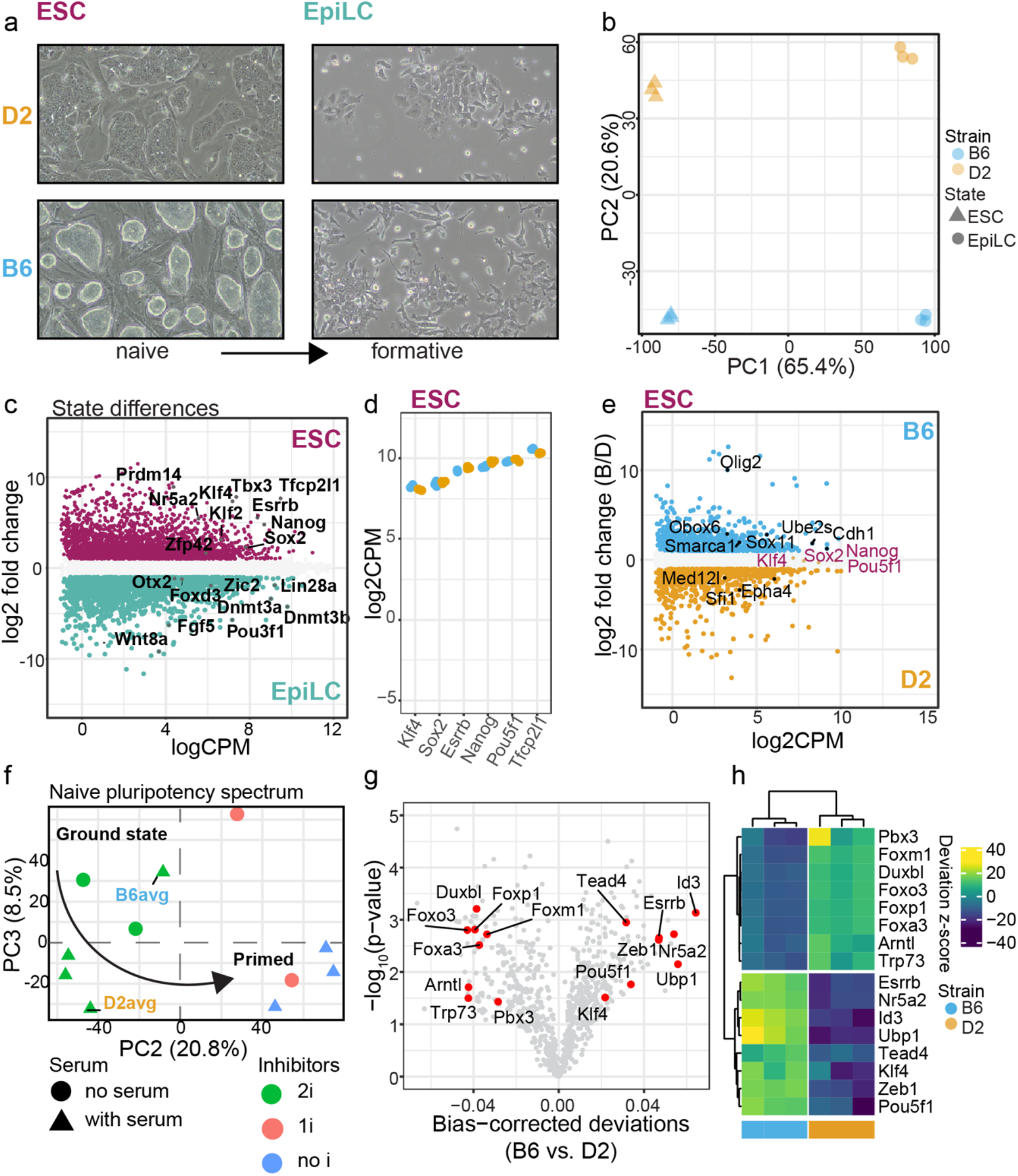
Genetic background influences position within a pluripotency spectrum through differences in chromatin accessibility. **a**, Phase contrast images of representative cultures for B6 and D2 strains maintained in conditions supporting naive pluripotency and after transitioning to formative EpiLCs. B6 ESC colonies adopt a classical ground state morphology, whereas D2 ESCs grew with a flat morphology and fewer cells per colony. Morphological differences persisted in EpiLCs. **b**, First two principal-components shows major source of variation in transcript abundance is cell state (PC1, 65.4%) and strain (PC2, 20.6%) for 3 biological replicates of B6 and D2. **c**, MA plot highlighting genes differentially expressed between state independent of strain (glm state term, false discovery rate-adjusted *P* value (FDR) < 0.05 and log_2_FC >1). Core pluripotent TFs highlighted for each state (maroon = naive ESCs, teal = formative EpiLCs). **d**, Scatterplot of expression for core naive TFs in ESCs. Equal high expression of naive TFs observed in both B6 and D2 ESCs. **e,** MA plot showing differentially regulated genes between strains within ESCs as determined by pairwise comparison (FDR < 0.05 and logFC >1). Core naive TFs are highlighted as not significantly differentially expressed between strains. Genes significantly different between strains within cell state with known roles in promoting naive pluripotency (B6 = *Obox6, Cdh1, Smarca1, Ube2s*; D2 = *Med12l, Sfi1*) as well as self-renewal in progenitor cell populations are highlighted (B6 = *Olig2, Sox11*; D2 = *Epha4*). **f**, PCA of full transcript abundance comparing average expression from B6 and D2 ESCs collected here to previously collected RNA-seq from Hackett et al. (2017). PC1 separated experimental origin, indicating batch effects between labs, and was not explored further. B6 expression was similar to that of ground state conditions which exclude serum; whereas D2 expression more closely resembled conditions that include serum. **g**, Volcano plot showing the mean difference in bias-corrected accessibility at TF motifs from ATAC-seq versus *P* value for that difference. Select TFs critical to pluripotency and differentiation are highlighted. **h**, Heatmap of motif deviations from 3 biological replicates of B6 and D2 ESCs for TFs highlighted in **g**.

Gene set enrichment analysis in ESCs identified strain-specific transcriptional programs. B6 enriched pathways included metabolism of amino acids and chromatin organization (**Extended Data Fig. 1d**). Derivatives of amino acid metabolism sustain nucleotide synthesis, required for rapid proliferation^39^ in ESCs, and serve as donors for histone modifications governing the pluripotent epigenetic landscape^40^. Additionally, DEGs were enriched for pathways representing different cell cycle phases with B6 enriched for genes regulating M-phase and D2 enriched for genes regulating G2/M transition. Cell cycle phase impacts exit from ground state pluripotency as an extended G2 phase delays differentiation^41^. D2 DEGs were paradoxically enriched both for up regulation of anterior-posterior pattern specification (**Extended Data Fig. 1e**), typically activated after exit from pluripotency, and WNT signaling which prevents exit from ground state^42,43^.

Next we compared the transcriptional state of our ESCs to that of published B6 ESCs cultured in nine conditions covering a range of pluripotency^28^. PCA of total transcript abundance reconstructed the reported spectrum of pluripotency and showed that B6 and D2 occupied different positions along this continuum (**Fig. 1f and Extended Data Fig. 1f**). PC2 distinguished ESCs grown in 2i and PC3 separated conditions containing serum. Along PC2, both B6 and D2 ESCs were transcriptionally similar to conditions that include 2i; however, variation in PC3 suggested that our B6 ESCs were more similar to cells grown in conditions that excluded serum, whereas D2 was transcriptionally similar to cells grown in conditions that included serum.

Since differences in the pluripotent spectrum are not explained by expression of naive TFs in ESCs, we measured variability in chromatin accessibility as an indicator of TF occupancy. Naïve TFs, including ESRRB, TFCP2L1, KLF4, POU5F1, and NR5A2, all had greater accessibility at their respective motifs in B6 compared to D2 (**Fig. 1g&h)**. Additionally, the motif for ZEB1, a TF important for neuronal differentaition^44^ is more accessible in B6 ESCs. In contrast, D2 ESCs showed greater accessibility of TRP73 motif, p53 family member known to instruct ESC differentiation towards mesoendoderm^45^. Given that transcript abundance of these TFs are similar between B6 and D2, this suggests the existence of unknown factors that regulate TF accessibility in *trans*. Combined, these data support that differential transcriptional programs, driven by genetic variation, place B6 ESCs in a more naïve position, while D2 ESCs are more primed to exit naive pluripotency, thus potentially impacting cell fate upon differentiation.

### Genetic background dictates activation of distinct biological processes upon exit from naïve pluripotency

Given observed differences between B6 and D2 in ESCs, we sought to understand how genetic background drives cell state transition. Our linear model identified 4,787 DEGs between cell states, 2,210 DEGs between genetic backgrounds and 2,083 genes with a significant genotype by state (GxS) interaction (FDR < 0.05 and log2FC >1, **Fig. 2a-d, Extended Data Fig. 2a-c, Extended Data Table 2**). This GxS interaction is exemplified by *Nr5a2* and *Sox1*, markers for pluripotency and neuronal progenitor cells respectively^46^. In ESCs, B6 and D2 show equally high expression of *Nr5a2,* yet upon exit from naïve state, D2 EpiLCs retain higher expression (**Fig. 2c**). In contrast, *Sox1* expression is equally low in B6 and D2 ESCs, increasing only in B6 EpiLCs (**Fig 2d**). To functionally characterize GxS interactions, DEGs were divided into modules using a hidden Markov model to classify unique expression paths, followed by filtering for genes with a significant GxS interaction (**Fig. 2e, Extended Data Table 3**). Example genes demonstrate each module’s unique trajectory upon transition from naive to formative pluripotency (**Extended Data Fig. 2d-o**). Importantly, different modules represent different biological processes. Nine modules were enriched for GO terms including neurogenesis, proliferation, and cell cycle regulation (**Fig. 2f and Extended Data Fig. 3a-e**). For example, module 4 genes (m4, n = 320 including *Nr5a2*) are initially higher expressed in B6 ESCs compared to D2; however, upon transitioning to EpiLCs D2 retains greater expression (**Fig. 2g**). When comparing module 4 genes in EpiLCs between strains, D2 shows enrichment by GSEA for genes specifically targeted for histone modifications in ESCs specifically, further suggesting D2 retainment of pluripotency programs in EpiLCs. Supporting the enrichment of M phase of cell cycle regulation in B6 ESCs, m9 (n = 10) was enriched for genes regulating mitotic cell cycle phase transition (**Fig. 2f&i**). Notably, module 5 (m5, n = 167, including *Sox1*) increased expression in B6 at a higher rate than D2 and is enriched for GO terms associated with neuronal development (**Fig. 2f&h**). These GxS interactions identify expression modules representing different paths during transition from naïve to formative pluripotency. Additionally, differential upregulation of lineage markers supports genetic biases in cell fate, with B6 primed towards neuronal lineages.

**Figure 2:**
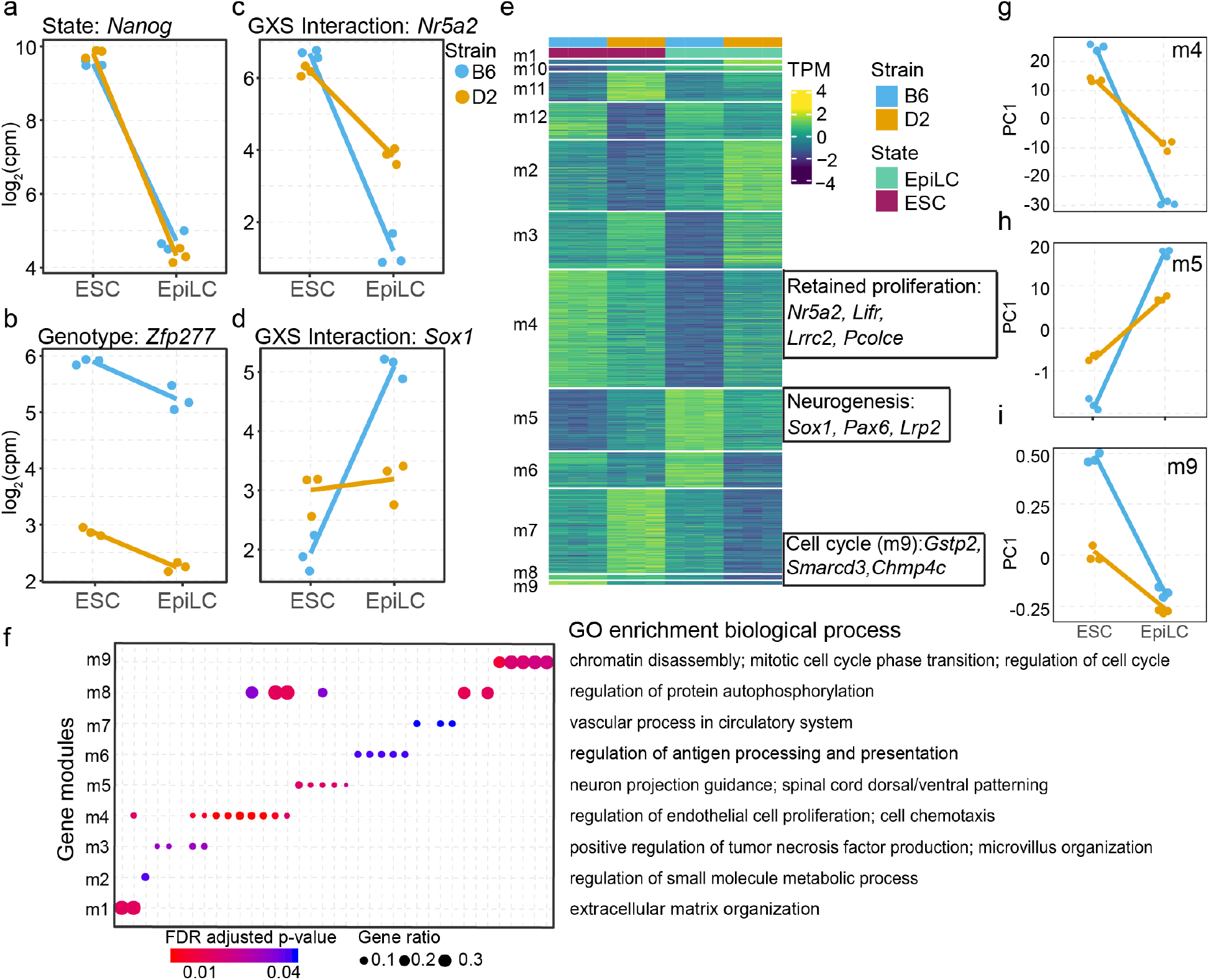
B6 EpiLCs are primed towards neuroectoderm lineage. **a-d**, Example expression patterns for select genes identified by applying a general linear model (glm) including state, genotype, and interaction terms. Dots represent individual biological replicates; lines highlight changes in mean values for replicates between each state (log_2_FC >1 and FDR < 0.05). **a**, State dependent expression is exemplified by *Nanog* which showed no difference between strains. **b**, Expression of *Zfp277* exemplifies strain-dependence being consistently higher in B6. **c & d**, A significant genotype x state (GXS) interaction was identified for *Nr5a2* (**c**, a marker for pluripotency) and *Sox1* (**d**, a marker for neuronal differentiation). **e**, Heatmap of transcript abundances for individual genes (rows) representing 12 expression modules detected by EBseqHMM filtered for genes with a significant GXS glm interaction. Genes with observed functional similarity within modules are highlighted. *Nr5a2*, *Lifr,* and other pluripotency genes shared the same expression path (m4) that decreases more significantly in B6, retained expression in D2, when ESCs are differentiated to EpiLCs. *Sox1, Pax6* and other neurogenesis genes shared an expression path (m5) that is significantly up regulated in B6 in EpiLCs. *Gstp2, Smarcd3* and other cell cycle regulation genes shared an expression path (m9) that is significantly up regulated in B6 ESCs. **f**, Gene ontology (GO) enrichment of biological processes (FDR adjusted *P* value < 0.05) identified for 9 out of 12 expression modules (m1-9). Each column represents a GO term. Highlighted terms are indicated to the right. GO terms associated with proliferation (m4), neurogenesis (m5) and cell cycle regulation (m9) indicate genetic control of exit from pluripotency and early lineage priming. **g-i**, Summarized differential module behavior between B6 and D2 upon transition from ESCs to EpiLCs represented by PC1 of transcript abundance for all genes within each module. **g**, D2 retained proliferation in EpiLCs. **h**, Up regulation of neurogenesis in B6 EpiLCs. **i**, Increased regulation of cell cycle M phase in B6 ESCs.

### Cellular systems genetics identifies *trans* regulation of chromatin accessibility and gene expression

To identify loci influencing differences between strains we took a cellular systems genetics approach by deriving ESCs from 33 individual BXD recombinant inbred mice, each representing a unique homozygous mosaic of the B6 and D2 founders^47^. RNA- and ATAC-seq were performed for both ESCs and EpiLCs. PCA of total RNA found that PC1 separated cell state (57.6% of total variance), while PC2-9 captured variance in genetic background (22.2% total variance, **Fig. 3a and Extended Data Fig. 4a-c**), supporting genetic governance over position within cell state.

**Figure 3:**
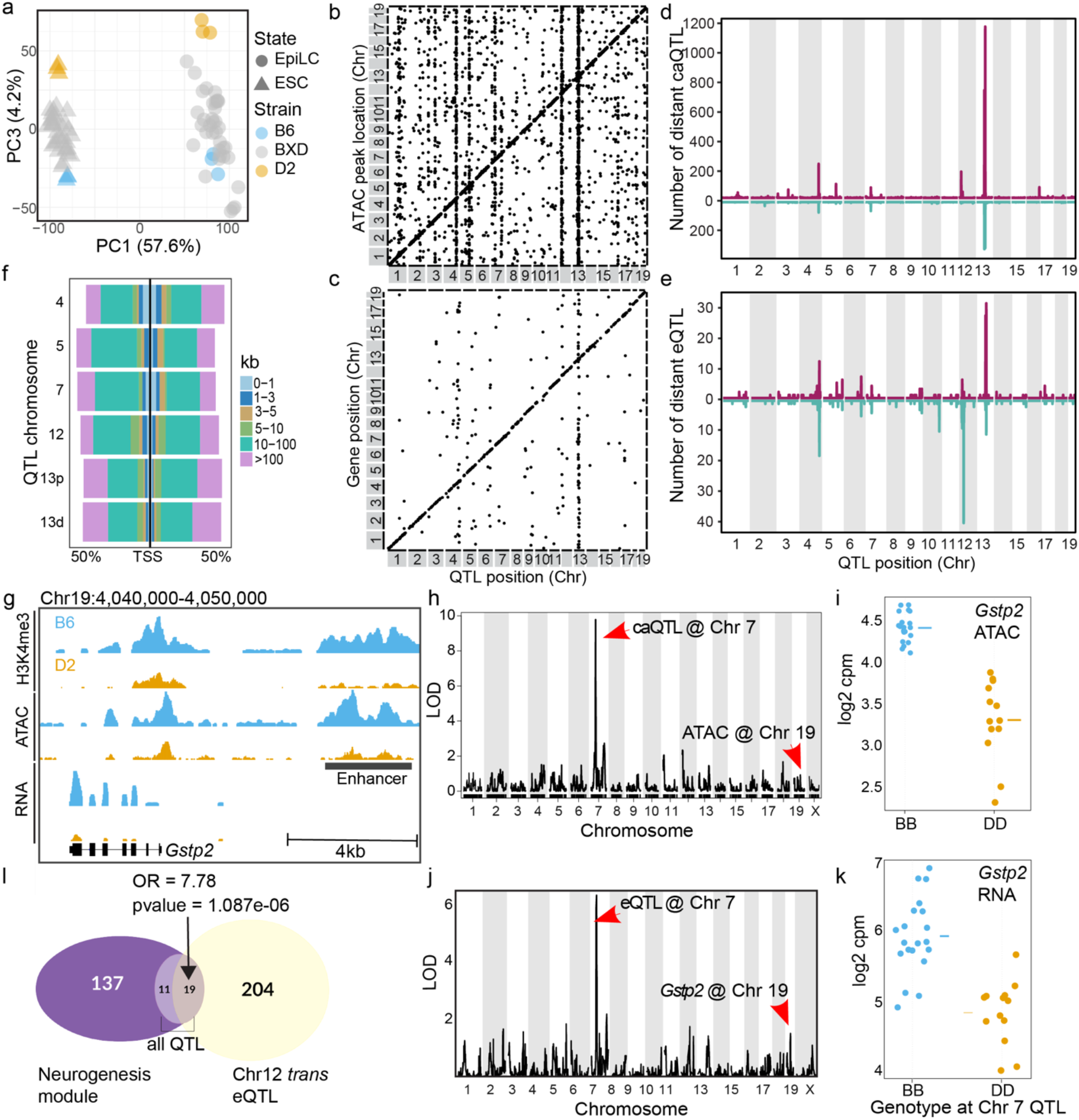
*trans*-acting quantitative trait loci co-regulate chromatin accessibility and gene expression in ESCs and EpiLCs. **a,** PCA of total transcript abundance in ESCs and EpiLCs from parental and 33 distinct recombinant inbred BXD strains generated from crosses between B6 and D2. PC1 captures variation between cell state (57.6%) and PC3 captures transcript variation within cell states. **b,** Scatterplot of genomic locations of individual chromatin accessibility (ca)QTL (x-axis, LOD > 5) versus location of ATAC peak being regulated (y-axis) for ESCs. Genetic effects that act locally (n = 3,973), i.e. in *cis*, fall on the diagonal line, while distal acting QTL (n = 13,767) lie off the diagonal. Several prominent distal QTL hotspots appear as vertical lines. **c**, Similar to b, plotting expression (e)QTL position versus gene location (n = 701 local-eQTL, n = 357 distal-eQTL). **d**, Number of distal caQTL in 1 Mb windows versus genomic location mapped in ESCs (maroon) and EpiLCs (teal). **e**, Similar to d comparing eQTL location and number between ESCs and EpiLCs. **f,** Distribution of caQTL hotspot targets in relation to nearest transcription start site in ESCs as percentage of total targets. **g**, Coverage profile for *Gstp2* locus on Chr 19 showing H3K4me3, ATAC, and RNA read depth. Putative upstream enhancer indicated with black bar. **h,** LOD score plot from QTL scan in ESCs for chromatin accessibility at the putative enhancer at *Gstp2*. Genetic variation at Chr 7 QTL distally regulates chromatin accessibility on Chr 19. **i**, Phenotype by genotype plot of the Chr 7 caQTL from h. A B6 haplotype at Chr 7 is associated with increased chromatin accessibility on Chr 19. **j&k**, Similar to h&i showing QTL mapping and haplotype effect for expression of *Gstp2* in ESCs. **l**, Euler diagram showing overlap between genes in the neurogenesis module (m5 from **fig. 2i)** and Chr 12 *trans* eQTL (LOD >4) in EpiLCs (enrichment compared to all eQTL in neurogenesis module, Fisher’s exact test – odds ratio (OR) = 7.78, *P* value = 1.087e-06).

Genetic mapping identified abundant local and distal chromatin accessibility (ca-) and gene expression (e-) quantitative trait loci (QTL) in both ESCs and EpiLCs (**Fig. 3b-e, Extended Data Tables 4-7**). Six prominent caQTL “hotspots” were identified in ESCs, most exhibiting shared distal co-regulation of chromatin accessibility and gene expression. Interestingly, these same QTL hotspots are also active in EpiLCs. Notably, EpiLC hotspots harbored a greater proportion of *trans* regulated gene targets accounting for 42% (84/200) of all distal-eQTL compared to 23% (82/357) in ESCs, whereas distal-caQTL were evenly regulated between states; 66.4% (778/1172) in EpiLC and 62.9% (2501/3973) in ESCs. Further, co-regulation was not confined within a cell state. The Chr 12 QTL hotspot regulates a greater proportion of putative regulatory elements in ESCs, but many more genes in EpiLCs, suggesting state-dependent *trans* regulation of molecular features (**Fig. 3d&e, Extended Data Fig. 4d-f**).

One causal molecular chain explaining shared regulation of distal chromatin accessibility and gene expression, is if a factor within the QTL regulates chromatin in *trans*, which then mediates local gene expression in *cis*. This would be evident by paired targets of ca- and eQTL mapping to the same locus. Indeed, most distal chromatin targets are 10-100 kb from the nearest promoter, suggesting putative regulatory elements that may act in *cis* (**Fig. 3f)**. *Gstp2*, located on Chr 19, is a member of module 9 associated with B6 enrichment of mitotic phase transition. Differential expression of *Gstp2* is correlated with differential chromatin accessibility at a nearby putative regulatory element (**Fig. 3g**), with B6 showing higher levels of both. QTL mapping in BXDs identified a single locus on Chr 7, with the B6 haplotype increasing both molecular features in *trans* (**Fig. 3h-k**). Extending these observations within cell type, about half (46.96%) of all genes regulated by a distal-eQTL hotspot were associated with a paired caQTL in ESCs; whereas, the majority of EpiLC QTL targets were not paired (**Extended Data Fig. 4g, Extended Data Table 8**). Interestingly, the Chr 12 hotspot was unique with a distinct bias towards paired ca- and eQTL across cell states (**Extended Data Fig. 4h**). While a single Chr 12 target gene matched a nearby caQTL in EpiLCs, 4 of the EpiLC gene targets were associated with chromatin targets in ESCs (**Extended Data Fig. 4h**). In EpiLCs, Chr 12 QTL target genes were significantly enriched, over all other eQTL, for GO terms associated with neuronal development programs (Fisher’s exact test, OR = 7.8, p-value = 1.087e-6; **Fig. 3l**). Similar to the observed neural bias in B6, the presence of a B6 haplotype in BXD-derived lines at Chr 12 lead to increased expression of the distal neuronal-associated genes. These data suggest that diffusible factors expressed from within the QTL hotspots variably regulate chromatin accessibility distally, ultimately leading to variation in transcript abundance of developmentally important genes.

### Occupancy of pluripotent TFs at regulatory elements is directed by caQTL

Given that expression levels of pluripotent TFs are similar, whereas accessibility of their binding motifs is variable, we hypothesized that causal genes underlying hotspot QTL regulate TF occupancy in *trans*. We calculated enrichment of overlap between publicly curated ChIP-seq datasets performed in mESCs with our distal hotspot caQTL intervals (**Fig. 4a, Extended Data Table 9**). While binding of TRIM28, a chromatin repressor^48^, showed the most significant overlap across all caQTL targets, pluripotent TFs, enhancer activators and chromatin remodelers were all enriched depending on the QTL. For example, targets of QTL on Chrs 4, 5, 7, and 12 are all enriched for POU5F1, P300, NANOG and SOX2.

**Figure 4:**
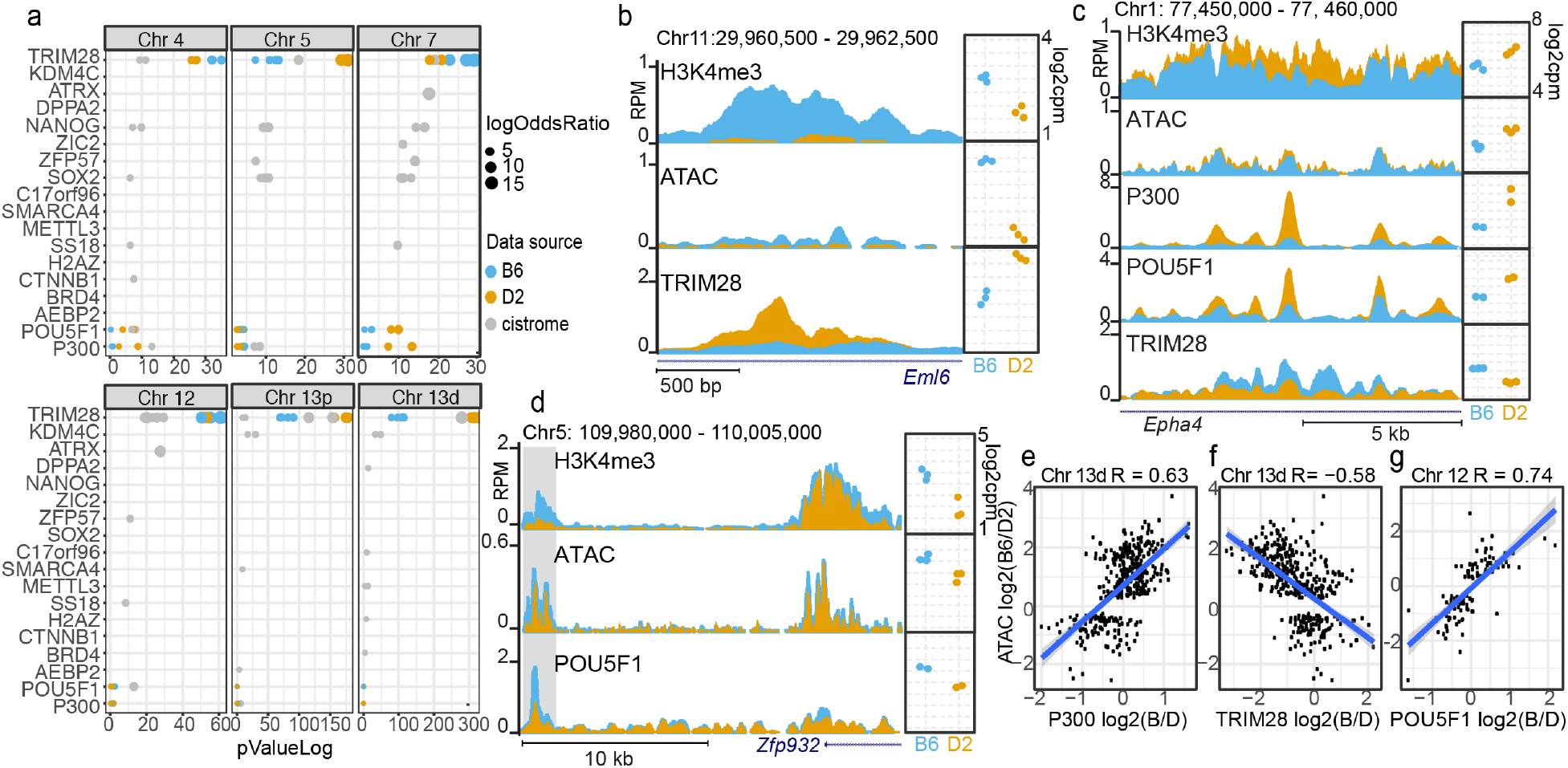
*trans*-QTL regulate chromatin binding of transcription factors required for pluripotency and differentiation. **a,** Enrichment of DNA-binding proteins at targets of hotspot caQTL in ESCs (Fisher’s exact test). Publicly available ChIP-seq datasets from Cistrome performed using ESCs (grey circles) were evaluated for significant overlap of QTL targets. Top 10 factors, ranked by maxrnk score from LOLA output, are plotted for each hotspot at q-value significance cutoff < 0.01. To validate a subset of these results ChIP-seq was performed for TRIM28, POU5F1, and P300 in B6 (blue) and D2 (orange) ESCs. Each circle represents biological replicate (B6 and D2) or independent ChIP-seq accession (Cistrome). **b-d,** Left – Coverage profiles for ChIP factor occupancy at representative putative regulatory elements targeted by QTL hotspots. Right – Scatterplots for ChIP factors comparing quantitative level of modification/binding for independent replicates between B6 and D2 ESCs. **b**, A putative regulatory locus on Chr 11 in an *Eml6* intron is under *trans*-regulation by the Chr 13d QTL. Greater occupancy of TRIM28 in D2 is inversely associated with increased chromatin accessibility and higher H3K4me3 signal in B6. **c**, A putative regulatory element in intron 3 of *Epha4* is controlled by the Chr 4 QTL. Increased chromatin accessibility (ATAC), H3K4me3 modification, occupancy of P300, and POU5f1 in D2 ESCs is inversely associated with higher TRIM28 binding in B6. **d**, A putative regulatory element on Chr 5 near *Zfp932*, targeted by Chr 12 *trans* caQTL, showed greater occupancy of POU5F1 and increased open chromatin compared to D2. **e-g**, Scatterplot of log_2_ fold change in ChIP signal versus change in chromatin accessibility (ATAC) between B6 and D2 ESCs for indicated caQTL. **e**, P300 occupancy at all Chr 13d caQTL targets was positively correlated with ATAC signal (Pearson’s R = 0.63). **f,** TRIM28 occupancy at the same targets of Chr 13d caQTL in *e* were negatively correlated with ATAC signal (Pearson’s R = −0.58). **g**, POU5F1 occupancy was positively correlated with ATAC at all Chr 12 caQTL targets (Pearson’s R = 0.74).

To validate binding of TFs at QTL targets and test for differential occupancy between strains, we performed TRIM28, P300 and POU5F1 ChIP-seq and combined with our ATAC-seq and H3K4me3 ChIP-seq data^49^. For example, a Chr 13d caQTL target in the intron of *Eml6* showed higher TRIM28 binding in D2 correlated with reduced chromatin accessibility and H3K4me3 level (**Fig. 4b**). Reciprocally, the distally regulated *Epha4* locus showed increased accessibility and gene expression when D2 at the QTL on Chr 4. In agreement, H3K4me3, P300 and POU5F1 show higher levels in D2, and conversely in B6, greater TRIM28 occupancy and less accessibility (**Fig. 4c**). Finally, at an example Chr 12 QTL target, increased POU5F1 occupancy in B6 coincided with greater open chromatin (**Fig. 4d**). These examples can be generalized to most QTL targets. Distal targets of Chr 13d showed positive correlation between differential P300 binding and increased open chromatin between strains (**Fig. 4e**), whereas differential TRIM28 binding is negatively correlated with chromatin accessibility at the same loci (**Fig. 4f**). Similarly, Chr 12 distal caQTL targets show a positive correlation between differential POU5F1 binding and chromatin accessibility (**Fig. 4g**). Generally, when one parental background had higher TRIM28 repressor binding, the other parent exhibited higher TF binding (**Fig. 4a** colored points, **Extended Data Table 10**). Together, these data show that *trans* regulation of chromatin in ESCs has a significant impact on regulatory elements bound by factors critical for establishment and maintenance of pluripotency.

### Genetic background biases differentiation propensity

To determine how genetic differences in pluripotency impacts cell fate, we developed an assay for spontaneous differentiation of embryoid bodies (EBs) and profiled these spheroids by single-cell RNA-seq (scRNA-seq) to determine cell composition. Cellular populations within EBs were highly reproducible among biological replicates (**Fig. 5a**, **Extended Data Fig. 5a-g**), permitting comparisons between strains. After integration of scRNA-seq across genetic backgrounds followed by unbiased clustering, 9 clusters could be annotated with identifiable cellular lineages (**Fig. 5b**, **Extended Data Table 11**). The remaining clusters were enriched for cellular activities representative of many cell types, such as mitotic cell cycle regulation or chromatin organization, precluding exact lineage identification. Importantly, different genetic backgrounds are biased towards different cell fates (**Fig. 5c&d**). For example, cells in cluster 9 represent primitive erythrocytes and are largely of D2 origin, whereas cells in cluster 5 express vascular endothelium marker genes and are largely B6. Cluster 4, showing equal proportions between strains, lies at the convergence of clusters 5 and 9 and represents yolk sac blood island cells (**Extended Data Fig. 5h**), common progenitors of primitive erythrocytes and vascular endothelium^50^. This suggests that genetic variation modulates trajectories through this developmental bifurcation. In addition to differences in cell fate within the same germ layer (i.e. mesoderm), EBs also identified differentiation bias to primary germ layers. Critically, definitive endoderm (cluster 7) was comprised mainly of D2 cells, whereas neuroectoderm (cluster 11) was comprised mainly of B6 cells.

**Figure 5.**
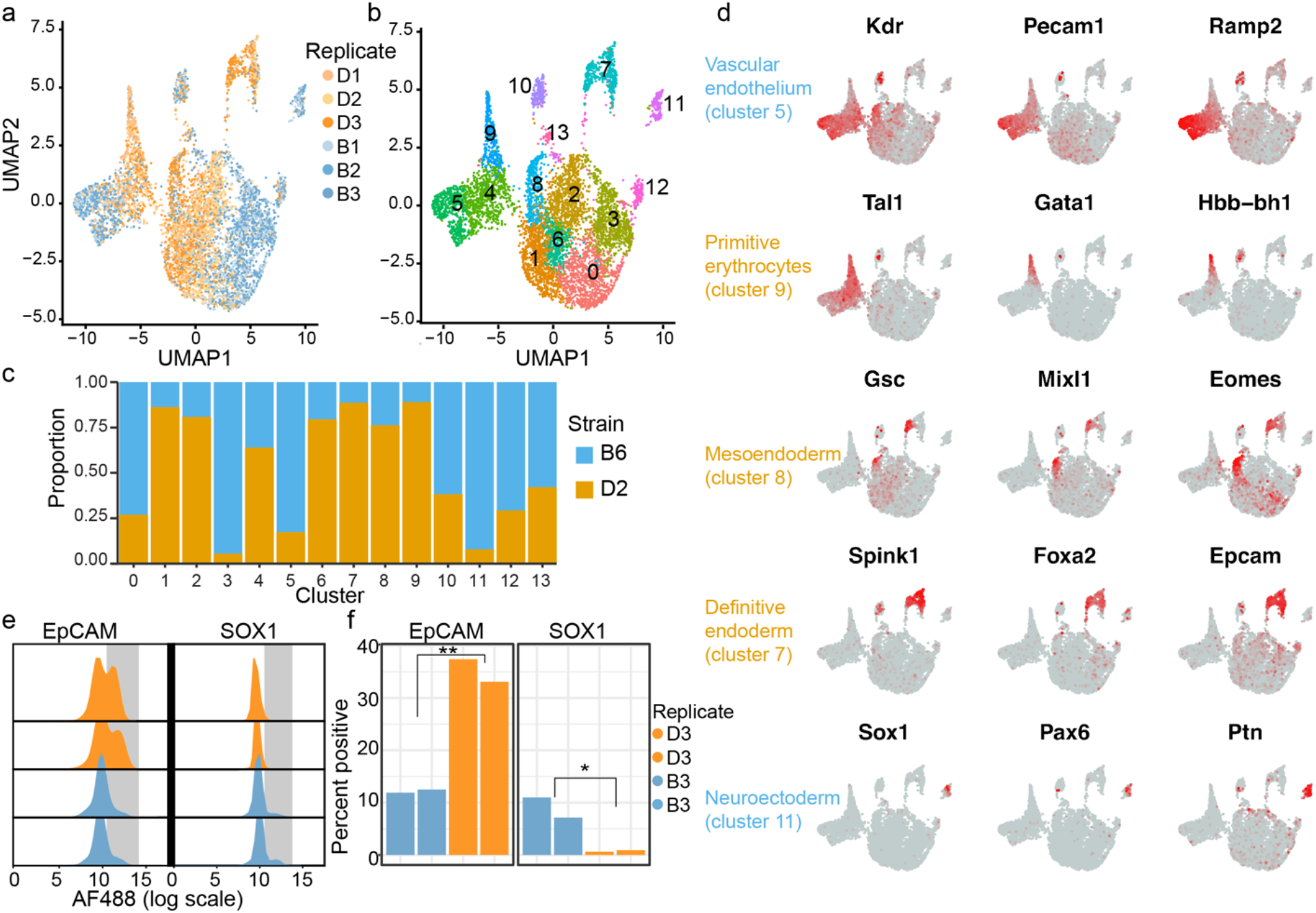
Embryoid bodies confirm genetic background influences differentiation propensity with B6 trajectory towards neuroectoderm. **a,** UMAP embeddings of transcriptional profiles from spontaneously differentiated embryoid bodies (EBs). Each point represents a cell colored by genetic background and shaded by biological replicate. **b**, Similar to *a* indicating 13 major cell populations based on unsupervised clustering. Each cell is colored based on cluster. **c,** Proportion of cells in a given cluster from *b* represented by either B6 or D2 genetic background. **d,** Feature plot of expression gradients for indicated gene overlaid on UMAP from a. Three lineage markers that help to distinguish cellular identity are shown for five cell clusters. **e**, Density plots show distribution of fluorescence from FACS analysis of protein abundance using antibodies representing lineage markers for definitive endoderm (left – EpCAM) and neuroectoderm (right – SOX1). Positive population of EB cells expressing lineage marker highlighted in grey. **f,** Bar chart showing percent positive cells from *e* gated for EpCAM+ population (left) or SOX1+ population (right). As predicted by scRNA-seq, more D2 EB cells express EpCAM compared to B6 and more B6 EB cells express SOX1 compared to D2 (test, ** pvalue < 0.01, * pvalue <0.05).

To validate transcriptional measurements of differentiation bias, FACS analysis was performed on protein abundance using antibodies against cell lineage markers. Compared to B6, a greater proportion of D2 EB cells expressed EpCAM, indicating definitive endoderm differentiation (12.2% vs. 35.25%, respectively, **Fig. 5e&f)**. Likewise, a greater proportion of cells from B6 EBs expressed SOX1 compared to D2 (9.07% vs. 0.76%, **Fig. 5e&f, Extended Data Fig. 5i**), indicating neuroectoderm bias. In summary, differentiation propensity is predominantly driven by genetic background. Importantly, B6 bias in neuroectoderm differentiation discovered in EBs is consistent with lineage priming of neuronal development evident in EpiLCs.

## Discussion

Here we provide a rationally designed genetic framework to critically assess genetic contribution to differentiation potential. Our data suggests that ESCs from B6 and D2 genetic backgrounds 1) reside at different locations along a pluripotent spectrum, 2) exhibit divergence in gene networks during transition from naïve to formative pluripotency, and 3) harbor at least 6 loci that distally co-regulate chromatin accessibility, transcription factor binding, and gene expression; all of which are consistent with outcomes that alter cell fate specification.

Modeling *in vivo* developmental progression *in vitro* has captured critical intermediate states between naïve and primed pluripotency^22^. Formulation of culture conditions permitting propagation of intermediate states, such as peri-implantation rosettes, has revealed aspects of pluripotent gene networks important for balancing maintenance of self-renewal with competency for differentiation^51,52^. Here, we show ESCs derived from B6 and D2 mice achieve occupancy of discrete states along the pluripotent spectrum. B6 ESCs acquire transcriptional profiles closer to ground state pluripotency, even in undefined media conditions, and upon transitioning to formative pluripotency rapidly dissolves the naïve program along with apparent priming towards neuroectoderm. In contrast, D2 ESCs exits pluripotency more slowly, as evidenced by stable expression of naïve TFs with little overt lineage bias.

Even though B6 and D2 ESCs occupy different positions along the pluripotent spectrum, they express similar levels of naïve markers, providing a conundrum as to where differences arise. To reconcile this, we adopted a multi-omics approach, measuring chromatin accessibility as a gauge for regulatory element activity. This approach showed that chromatin accessibility at pluripotent TF motifs are higher in B6 compared to D2, indicating other *trans* acting chromatin regulation may determine pluripotent spectrum. We identified 6 QTL hotspots that alter binding of naïve TFs in *trans*, and ultimately differentially regulate proximal genes pertinent to cell fate. Our study additionally suggests chromatin priming in ESCs influences gene expression in EpiLCs, for example instructing commitment to neuroectoderm. The fact that a greater percentage of ca- and eQTL are paired in ESCs but not EpiLCs could suggest that the primary effect of QTL on influencing chromatin accessibility resides in ESCs, ultimately leading to secondary or tertiary gene expression changes after differentiation. In EpiLCs, these downstream expression changes would still map back to the same distal locus identified in ESCs but may no longer be accompanied by local changes in chromatin directly. This finding supports recent observations of chromatin priming preceding cell-fate decisions during gastrulation^53^ and highlights the potential for early events in development to have differential cascading effects on adult phenotypes^54^.

Several lines of evidence support QTL discovered in this study are of significant developmental importance, best exemplified by the B6 trajectory towards neuroectoderm. First, genotype-by-state interactions during transition to formative pluripotency uncovered strain divergence during exit from naïve pluripotency and were responsible for installing competency of lineage specification with a predisposition to neuroectoderm in B6. Second, genetic mapping in BXDs identified a locus on Chr 12 regulating chromatin accessibility distally in ESCs thereby priming a regulatory landscape directing neuronal development in cells harboring a B6 haplotype at distal Chr 12 QTL. Finally, spontaneous differentiation to EBs confirmed B6 trajectory towards neuroectoderm. Aside from roles in governing cell state transitions described here, QTL at the same positions on Chrs 4, 12, and 13 have all been implicated in a variety of developmental and disease phenotypes such as limb abnormalities^55^, cleft palate^56^, lupus^57^, as well as association with differences in priming sites of meiotic recombination^49^ and stabilization of ground state pluripotency^14^. Notably, a single KZFP contained within our Chr 13 hotspot QTL interval was shown to be causal in the progression of a lupus phenotype^57^. And while the other studies largely have unknown causal factors implicated in reported phenotypes, overlapping QTL harbor clusters of newly emergent KZFPs in the murine genome^58,59^. This provides exciting future work into assigning causality to a rapidly evolving gene family whose divergence in different strain backgrounds may account for evolution of regulatory function that shapes development and disease^60^.

In the absence of inductive signals to direct differentiation, neuralization has been described as the ‘default’ development path^61^. *In vivo* inhibition of neuronal programs is due to extrinsic signaling originating from extraembryonic tissue, of which ESC cultures are typically devoid^62^. Historically, the study of differentiation *in vitro* has been limited to ESCs derived from B6 or 129 mouse strains^53,63^. Our study, however, suggests this neuronal default paradigm may not be universal to all genetic backgrounds, supporting recent findings showing mESC strain differences in neuralization capacity^15^. It is, therefore, paramount to assess causal factors underlying the Chr 12 QTL which are likely to play a role in capacitation for multi-lineage differentiation.

In total, this work demonstrates that ESCs derived from genetically diverse strains do not share equal developmental potential *in vitro*. Recent experiments have shown that differences in cell fate choice during development may be critical in predisposing individuals to complex diseases due to underlying differences in cell type composition^64^. Clearly, further investigation into genetic governance of differentiation is needed to understand its potential role in complex traits.

## Methods

### Derivation of mouse embryonic stem cells

Mouse embryonic stem cells were derived and maintained in conditions previously reported^6^. All mice were obtained from The Jackson Laboratory (Bar Harbor, ME) including C57BL/6J (stock number 000664), DBA/2J (stock number 100006), and BXD recombinant inbred lines (see **Extended Data Table 12**). All animal experiments were approved by the Animal Care and Use Committee of The Jackson Laboratory (summary #04008).

### Cell Culture

#### Naive pluripotency supported in ESCs

A vial containing 3 million ESCs (P4–P6) were thawed onto a 60 mm tissue-treated culture dish seeded with irradiated mouse embryonic fibroblasts (MEFs) as feeders in DMEM high glucose base medium supplemented with 15% fetal bovine serum (FBS, Lonza cat. no. 14-501F lot no. 0000217266), 1X Pen/Strep, 2mM GlutaMAX, 1 mM sodium pyruvate, 0.1 mM MEM NEAA, 0.1mM 2-Mercaptoethanol, 10^3^ IU LIF, 1 μM PD0325901, and 3 μM CHIR99021. ESCs were expanded for 2-3 days to reach ~70% confluency for molecular and differentiation assays.

#### Differentiation to formative pluripotency in EpiLCs

EpiLCs were grown from mESCs as previously described^21,37^. Confluent ESCs were washed with 1X PBS, and trypsizined (0.05%) to form single cell suspension, filtered through a 40uM mesh, and MEFs were excluded by settling on 60 mm dishes coated with 0.01% gelatin for 30 minutes. ESCs were seeded at a density of 200,000 cells per 60 mm dish coated with 5ug/mL fibronectin in medium containing N2B27 supplemented with 2 mM GlutaMAX, 1 mM sodium pyruvate, 1X Pen/Strep, 0.1 mM MEM NEAA, 0.1mM 2-Mercaptoethanol, 1% KOSR, and 12ng/mL bFGF. Cells were transitioned to not exceed 48 hours. Cells were gently dissociated using TrypLE (Thermo Fisher Scientific, cat. no. 12-605-010) for subsequent molecular and differentiation assays.

#### Spontaneous differentiation to EBs

To form EBs, EpiLCs washed with 1X PBS and dissociated with TrypLE and resuspended in EB medium containing DMEM high glucose base supplemented with 15% FBS, 2mM GlutaMAX, 0.1mM MEM NEAA, 1mM sodium pyruvate, 1X Pen/Strep, 0.1mM 2-Mercaptoethanol, and 12ng/mL bFGF at a seeding density of 750 cells/EB. Uniform and reproducible EBs were achieved using AggreWell 400 plates (STEMCELL Technologies, cat. no. 34415) containing 1200 microwells. Each well contains 2mL of single cell solution of 900,000 cells per well (750 cells/EB). Preparation of AggreWell 400 was followed per manufacturer’s instructions, with modification to only use middle 8 wells to ensure reproducibility of cultures. Cells were allowed to aggregate in microwells for 48 hours undisturbed. After 48 hours, each well was filtered through a 40um mesh to discard any unincorporated cells and collect newly formed EBs. Filter was then inverted and EB medium was passed through filter to collect EBs into 100mm Corning Ultra-low attachment culture dish (Millipore Sigma, cat. no. CLS3262). Dishes were placed on BellyButton rotator and left undisturbed for another 48 hours to allow EBs to further develop while limiting aggregation of individual EBs within culture. After 48 hours, EBs were filtered through a 40uM mesh to discard unincorporated cells. Single cell suspensions were achieved by incubating EBs in 0.25% trypsin for 2 minutes at 37°C followed by manual disaggregation with an Eppendorf P1000 manual pipette. Trypsin was inactivated by resuspending cells in PBS supplemented with 10% FBS and the cell suspension was filtered through a 40uM mesh. Cell suspensions were washed twice with PBS before proceeding with processing for scRNA-seq or FACS analysis.

### RNA-seq sample prep and sequencing

For mESC RNA collection, naive culture was trypsinized and MEFs were removed by plating single cell suspension onto gelatin-coated dishes and allowing MEFs to settle for 30 minutes. One million enriched mESCs were lysed and RNA extracted using the RNeasy (QIAGEN) RNA extraction kit. EpiLCs were harvested using TypLE and 1 million cells were used for RNA extraction. RNA concentration and quality were assessed using the Nanodrop 2000 spectrophotemeter (Thermo Scientific) and the RNA Total RNA Nano assay (Agilent Technologies). Libraries were constructed using the KAPA mRNA HyperPrep Kit (KAPA Biosystems), according to the manufacturer’s instructions. Library quality and concentration was checked using the D5000 Screen Tape (Agilent Technologies) and quantitative PCR (KAPA Biosystems).

### ATAC-seq and ChIP-seq sample prep and sequencing

To measure chromatin accessibility in ESCs and EpiLCs, cells were harvested as described above. For parental ESCs and EpiLCs along with BXD ESCs, the OMNI-ATAC^65^ protocol was followed using 100,000 cells. For BXD EpiLCs, the FAST-ATAC^66^ protocol was followed using 100,000 cells. Libraries were amplified for a total of 8-10 cycles and DNA purified using AMPure XP beads (Beckman Coulter). The quality and concentration of ATAC-seq libraries were evaluated using the Bioanalyzer DNA High Sensitivity Assay (Agilent Technologies) and quantitative PCR (KAPA Biosystems).

For each biological replicate ESC ChIP-seq library, 10 million cells were harvested and fixed as previously described^14^. Cell lysis, chromatin fragmentation, dialysis, and immunoprecipitation was performed as described^67^. Immunoprecipitation was performed using antibodies against P300 (12ul, Bethyl A300-358A), POU5F1 (20ul, Cell Signaling Tech Oct-4A rabbit mAb, 5677S), and TRIM28 (10ul, Abcam 201C, ab22553). ChIP-seq libraries were constructed using the KAPA HyperPrep Kit (Roche Sequencing and Life Science). Quality and concentration of libraries were assessed using the High Sensitivity D5000 ScreenTape (Agilent Technologies) and KAPA Library Quantification Kit (Roche Sequencing and Life Sciences).

### Single cell RNA-seq sample prep and sequencing

Three biological replicate EBs, starting from independently derived ESCs, were grown for both B6 and D2 parental strains. Additionally, starting from just one of the derived D2 ESC lines, EBs were grown in triplicate to represent technical replicates within cell line. Finally, 3 aliquots of disaggregated cells from one of the D2 EB cultures was sub-sampled to determine reproducible representation cell populations within EBs as they represent heterogenous organoids. The MULTIseq protocol^68^ was used to pool single-cell suspensions from EBs across samples to load onto one 10X Chromium lane to reduce batch effects. Each of the 10 samples outlined above were labeled with a unique lipid modified oligo following the published protocol using 500,000 cells per sample. After labeling, cells were counted and adjusted so that the final pool represented 3,000 cells per sample. Pooled cells were washed once in PBS and resuspended in PBS containing 1% BSA supplemented with 1% FBS. A single 10X Chromium microfluidic lane was super-load with 40,000 cells with the goal of obtaining approximately 1,000 barcoded cells per sample after sequencing, demultiplexing, and filtering. Single cell capture, barcoding, and library preparation were performed using the 10X Chromium version 3 chemistry, according to the manufacturer’s protocol (#CG00183). cDNA and barcode libraries were checked for quality on an Agilent 4200 TapeStation, quantified by KAPA qPCR, and sequenced on a single lane (95% transcriptome, 5% barcode) on Novaseq6000 (Illumina) to an average depth of 100,000 reads per cell.

### Fluorescent assisted cell sorting

Spontaneously derived embryoid bodies were independently grown from parental ESC lines to represent technical replicates. Single cell suspensions were fixed with 4% paraformaldehyde at a concentration of 1 million cells/mL for 15 minutes at room temperature. Cells were washed with PBS/1% FBS twice before antibody labeling. Cells were blocked for 15 minutes in either PBS/1%FBS (EpCAM) or PermWash (SOX1) prior to incubation with primary antibody (anti-EpCAM at 1/10k, abcam ab71916; anti-SOX1 at 1/300, R&D Systems AF3369) for 45 minutes at room temperature. After washing three times for 5 minutes each in PBS/1% FBS (PermWash for SOX1 samples) cells were incubated with secondary antibody for 45 minutes at room temperature. After washing three times for 5 minutes each in PBS/1% FBS (PermWash for SOX1 samples), cells were analyzed on Attune NXT Analyzer (Thermo Fisher). Cells were gated using FlowJo™ Software and density of cell populations were visualized using flowVis package in R (https://bioconductor.org/packages/flowViz/).

### Data Analysis

Statistical analysis was performed using R version 3.4.1 for qtl mapping, v.3.5.1 for normalization and transformation of inputs for downstream analysis, and v.3.6.2 for visualization and analysis (R Core Team, 2018).

#### ATAC-, ChIP-, and bulk RNA-seq data processing

Bulk parental and BXD ESC and EpiLC fastq files were aligned using bowtie^69^ to their respective strain-specific transcriptomes (Ensembl release 84) and transcripts were quantified at gene-level abundances using EMASE^70^. Read counts were filtered for lowly expressed genes (3cpm in at least 3 samples), TMM normalized using edgeR^71^ and log_2_ transformed for differential analysis and QTL mapping.

Illumina adaptors were trimmed from ATAC and ChIP reads using Trimmomatic (version 0.33) and then aligned to mouse reference genome (mm10, modified to incorporate known D2 variants reported in Mouse Genomes Project Database, REL-1505^72,73^) using hisat2^74^. Duplicate reads were removed using Picard Tools (“Picard Toolkit.” 2019. Broad Institute, GitHub Repository. http://broadinstitute.github.io/picard/; Broad Institute) and peaks were called for each sample using MACS^75^ (version 1.4.2, p = e^−5^). A comprehensive set of open chromatin locations (i.e. peakome) was generated by combining all ATAC peaks identified across all 33 BXD samples and peaks overlapping by 1 bp merged using bedtools. A similar peakome strategy was used for ChIP samples. ATAC and ChIP read counts were collected using bedtools^76^ multicov within respective peakome intervals. Read matrices were TMM normalized and log_2_ transformed for QTL mapping of ATAC samples in BXDs and downstream differential analysis of chromatin accessibility and ChIP factor occupancy between parental strains.

#### caQTL and eQTL mapping in ESCs and EpiLCs

Normalized and transformed read counts were used as input for QTL mapping for both transcript abundance and chromatin accessibility. QTL were mapped using the ‘scan1’ function in R/qtl2^77^ using a linear mixed model to account for kinship. Processing of BXD samples was included in the model as a covariate to account for batch effects (**Extended Data Table 13)**. To assess genome-wide significance in ESC RNA-seq samples, we calculated empirical p-values using a permutation strategy (1000 permutations) and applied a multiple testing correction to obtain FDR values (Benhamini-Hochberg adjusted p-value). Figure 3 reports significant eQTL with LOD cutoff > 5 and caQTL with LOD cutoff > 5 (**Extended Data Tables 4-7)**.

Distribution of hotspot caQTL from nearest TSS was determined using ChIPseeker^78^. Target genes of caQTL were assigned using GREAT^79^ and further filtered for genes with eQTL mapped to same genomic interval as paired caQTL (within and across cell state.

#### Differential analysis of gene expression

Differentially expressed transcripts between parental ESCs were determined using pairwise comparisons in edgeR performing quasi-likelihood F-test (FDR<0.05, log_2_FC>1). To determine sets of genes enriched in functional programs, we used significantly differentially expressed transcripts between strains (FDR<0.05) as in input in gene set enrichment analysis (GSEA)^80,81^. Strain specific enriched programs were identified using gene sets curated with MsigDB^80,82,83^ and considered significant at FDR < 25% (weighted scoring and 1000 permutations on phenotypes) and suggestive at nominal pvalue < 0.05 (1000 permutations on gene set).

Placement of parental ESCs along naive pluripotency spectrum was determined using previously published RNA-seq data from ESCs grown in 9 different culture conditions^28^. Raw fastq files were accessed in GEO (Accession: GSE98517) and uniformly processed to match our RNA-seq data as described above. Principle components were determined using prcomp function in R on TMM normalized and log_2_ transformed RNA read counts from the whole transcriptome. Gene contribution to separation of samples along PCs was achieved using factoextra package in R (https://rpkgs.datanovia.com/factoextra/index.html).

To determine differentially expressed transcripts dependent on cell state, strain, and a significant interaction between state and strain a general linear model (glm; Y = B_0_ + B_1_X_1_ + B_2_X_2_ + B_3_X_1_X_2_ where X_1_=state, X_2_=strain, and X_1_X_2_=interaction term) was applied to normalized RNA read counts from parental ESC and EpiLC samples. Unique expression paths of genes changing similarly across cell were identified using EBseqHMM^84^ (confidence cutoff = 0.5). Modules were defined by filtering EBseq paths for genes with a significant state by strain interaction (X_1_X_2_) from the glm. Principle component analysis was performed separately on each module using the expression of all genes within the module to summarizing the change in expression as an Eigen value. Gene modules and significantly differentially expressed genes identified in glm were visualized using ComplexHeatmap package in R^85^. Gene modules with enriched GO terms were determined using clusterProfiler in R^86^ (adjusted pvalue < 0.05).

#### Differential analysis of chromatin accessibility and ChIP factor occupancy

Differential accessibility of transcription factor motifs between parental ESCs as well as differences detected across cell states were determined using chromVar^87^. Unsupervised hierarchical clustering was performed on GC bias-corrected deviation z-scores.

To identify differential occupancy of factors at caQTL in ESCs we tested for overlap of Cistrome^88,89^ curated ChIP-seq datasets (mouse factor data downloaded 1/18/19, we extracted mouse ESCs ChIP-seq for use in this study representing 1,763 ChIP-seq experiments) along with ChIP-seq performed in this study (TRIM28, P300, POU5F1) using LOLA^90^ package in R. Regions were defined as the collection of chromatin targets intervals for each hotspot caQTL, the universe was set as the total ATAC-seq peakome and LOLA results were filtered by maxRnk (combined score for p-value and odds ratio). Allele dependent ChIP factor binding (P300, TRIM28, POU5F1) correlated with chromatin accessibility at hotspot caQTL was determined using Pearson’s correlation.

#### Analysis and annotation of single-cell RNA-seq

CellRanger was used to align scRNA-seq fastq files and identify individual cell barcodes from 10X Genomics data retrieving 20,134 individual cell transcriptomes. Individual cells were further classified into Multiseq lipid hash samples using demultiplex R package (https://github.com/chris-mcginnis-ucsf/MULTI-seq) resulting in 2,150 negative cells (no hash barcode), 3,022 doublets (identified as two hash barcodes) and 14,962 unique cells accounting for ~ 1,000 cells/sample (**Extended Data Table 14)**. Clustering and analysis of scRNA-seq data was performed using Seurat^91^. Cells with greater than 10% of reads from mitochondria genes were removed. Preprocess, normalization and dimensional reduction largely followed default Seurat settings including FindVariableFeatures (nfeatures=2000), RunPCA (npcs=100) with 20 principle components selected for clustering. Because of the large observed differences in clustering between B6 and D2 cells we used Harmony^92^ to mitigate the impact of cell strain background on clustering with the following settings (theta=1, dims.use=1:20, max.iter.harmony=100). Increasing theta resulted in forced clustering between cells with little biological meaning (data not shown), while not improving integration between genetic background. Therefore theta = 1 was chosen to maximize strain integration while preserving cell identity.

Unique and differentially expressed genes in cell clusters were generated from Seurat and subsequent gene lists were used to annotate cell clusters using MouseMine^93^ along with literature searches of top 5 unique and highly expressed genes in each cluster. MouseMine is a public database supporting gene set queries to discover functional relatedness to a biological process, interactions with other genes, as well as assess spatial expression within relevant anatomical regions of the developing mouse embryo. Nine cell clusters were annotated as a unique cell lineage and four clusters were enriched for GO terms describing cellular activities with no enrichment in anatomical structures or lineage markers.

Differentiation trajectory of cells in cluster 4 branching to clusters 5 and 9 was inferred using Slingshot^94^. Cells in cluster 4 were selected as starting cluster based on prior knowledge of gene markers indicating yolk sac blood island cells.

## Data Availability

## Acknowledgements

We would like to thank all members of the Baker laboratory for comments and discussion. Additionally, we would like to thank Dr. Martin Pera for constructive feedback on the project and reviewing the manuscript as well as Dr. Martin Ringwald for assisting with approaches to annotate developmental stages of single cell clusters. We recognize contributions from The Jackson Laboratory Genome Technologies Services and Flow Cytometry Services for technical assistance and consultation. The Jackson Laboratory scientific services are supported in part through the National Institutes of Health Cancer Core grant CA34196. The Special Mouse Strain Resource, NIH OD011102 (L.G.R.) provided the BXD recombinant inbred strains used for mESC derivation. This work is supported by The Jackson Laboratory Directors Innovation Fund to S.C.M., L.G.R., D.A.S., and C.L.B., C.B was supported by T32HD007065, and R35 GM133724 to C.L.B.

## Author Contributions

Conceptualization, C.L.B, C.B.; Methodology, C.L.B., C.B.; Resources, C.L.B., L.G.R.; Investigation, C.B., C.S., H.J.F., A.C.; Data Curation and Formal Analysis, C.B., C.L.B., C.S., D.A.S; Visualization, C.B., C.L.B., C.S.; Writing, C.B., C.L.B.; Revision and edits, C.L.B., D.A.S., L.G.R., S.C.M.; Supervision, C.L.B.; Funding Acquisition, C.L.B., L.G.R., S.C.M., D.A.S.

---

**Extended Data Figure 1:**
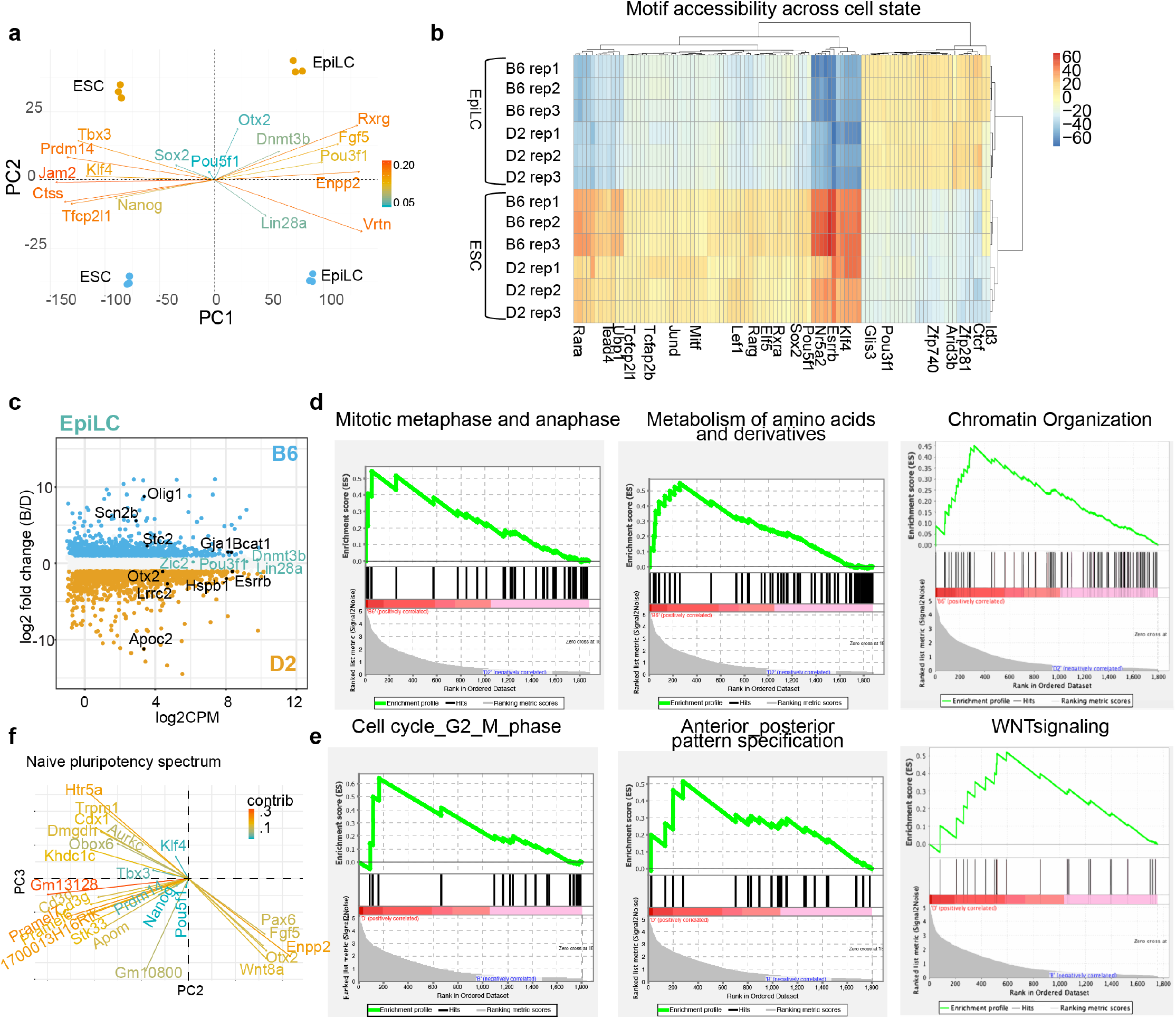
Expression of core naive TFs does not explain strain-specific transcriptional programs in ESCs. **a**, PCA bi-plot showing individual gene contributions to the first two principle components. Transcript abundance measured by bulk RNAseq from 3 biological replicates for each strain in ESCs and EpiLCs. Core naive and primed marker genes are indicated, drive separation between cell states, and are largely not differentially expressed between strains. As previously described (Buecker, 2014), *Pou5f1* expression is unchanged across cell states. **b**, Heatmap of bias-corrected deviation z-scores of chromatin accessibility at TF motifs (n = 3 biological replicates of ESCs and EpiLCs from both B6 and D2 strains). **c**, MA plot highlighting genes differentially expressed between strain within EpiLCsas determined by pairwise comparison (FDR < 0/05, log_2_FC > 1). Core primed TFs are highlighted as not significantly differentially expressed between strains. **d**, GSEA for functional programs curated within MsigDB using DEGs between B6 and D2 ESCs. Upregulated genes in B6 ESCs were enriched for GO terms in biological processes associated with self-renewal and pluripotency such as regulation of mitotic metaphase and anaphase and metabolism of amino acids and derivatives (significant enrichment in phenotype using 1000 permutations, FDR < 25%) as well as regulation of chromatin organization (significant for single gene set, nominal pvalue < 0.05). **e**, Similar to d showing pathways up regulated in D2. D2 ESCs were associated with regulation of G2/M phase transition and anterior-posterior pattern specification (FDR < 25%) in addition to self-renewing programs such as WNT signaling (nominal pvalue <0.05). **f,** PCA bi-plot indicating selected gene weights contributing to occupancy of cells along naive pluripotency spectrum from Figure 1f. Core naive TFs (such as *Klf4, Tbx3, Nanog, Pou5f1*, and *Prdm14*) do not contribute to separation of cells grown in different culture conditions or differential occupancy of pluripotency continuum by B6 and D2 ESCs. Cells grown in conditions lacking 2i have higher expression of primed TFs (i.e. *Fgf5, Otx2, Wnt8a*). While, B6 and D2 both associate with conditions that include 2i, D2 is separated from B6 ESCs by expressing transcription profiles similar to conditions that include serum. Genes weights selected are in top 100 genes contributing to separation along PC2 and PC3; only genes driving separation between B6 and D2 along PC3 are plotted at threshold of 0.15 contribution.

**Extended Data Figure 2:**
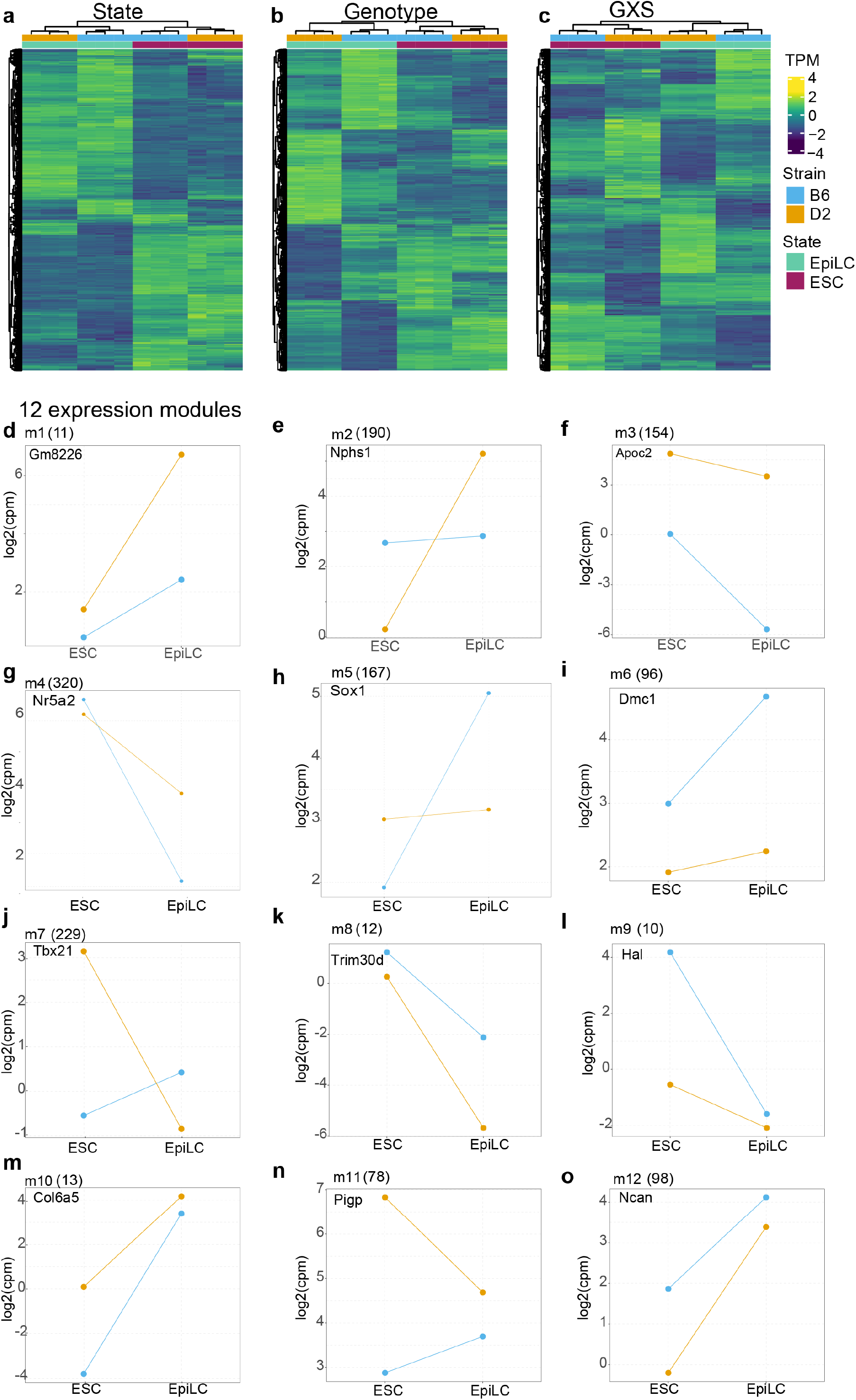
Genetic background induces distinct expression pattern changes across cell state transition. **a-c**, Heatmaps showing expression patterns for genes detected with a significant difference based on general linearized model (glm, state + strain + stain x strain). Hierarchical clustering of samples (columns) results in proper association of cell states and biological replicates within genotype. **a**, Genes with significant difference between cellular states independent of genetic background (n = 10,380 at FDR < 0.05). **b**, Genes with significant difference between strains independent of state (n = 7,359 at FDR < 0.05). **c**, Genes with significant interaction between state and strain (n = 4,912 at FDR < 0.05). **d-o**, Candidate genes expression patterns between genotype and state represent expression modules m1-12. Modules resulted from intersection between EBseqHMM defined expression paths and genes with a significant GxS interaction. Number of genes within each module indicated at top in parenthesis.

**Extended Data Figure 3:**
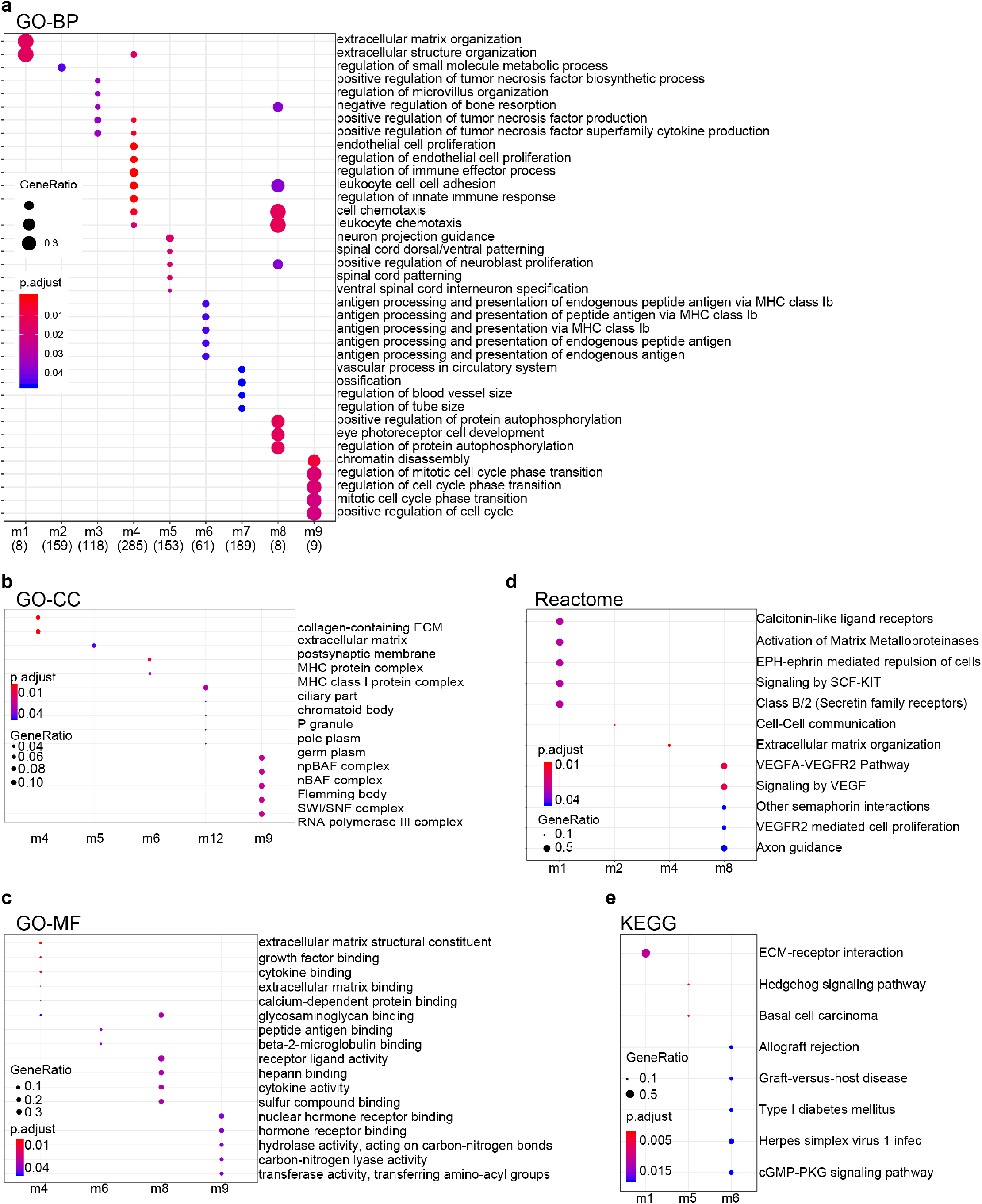
Discrete biological functions observed for each gene module. **a-e**, Significantly enriched GO terms indicating gene ratio (size of point; overlap of input with gene set/overlap of input with collection) and FDR adjusted P value (color) for GxS modules (m1-m12). GO enrichment associated with biological processes (**a**), cellular components (**b**), molecular function (**c**), as well as pathways curated in Reactome (**d**) and KEGG (**e**) are distinct for each expression module.

**Extended Data Figure 4:**
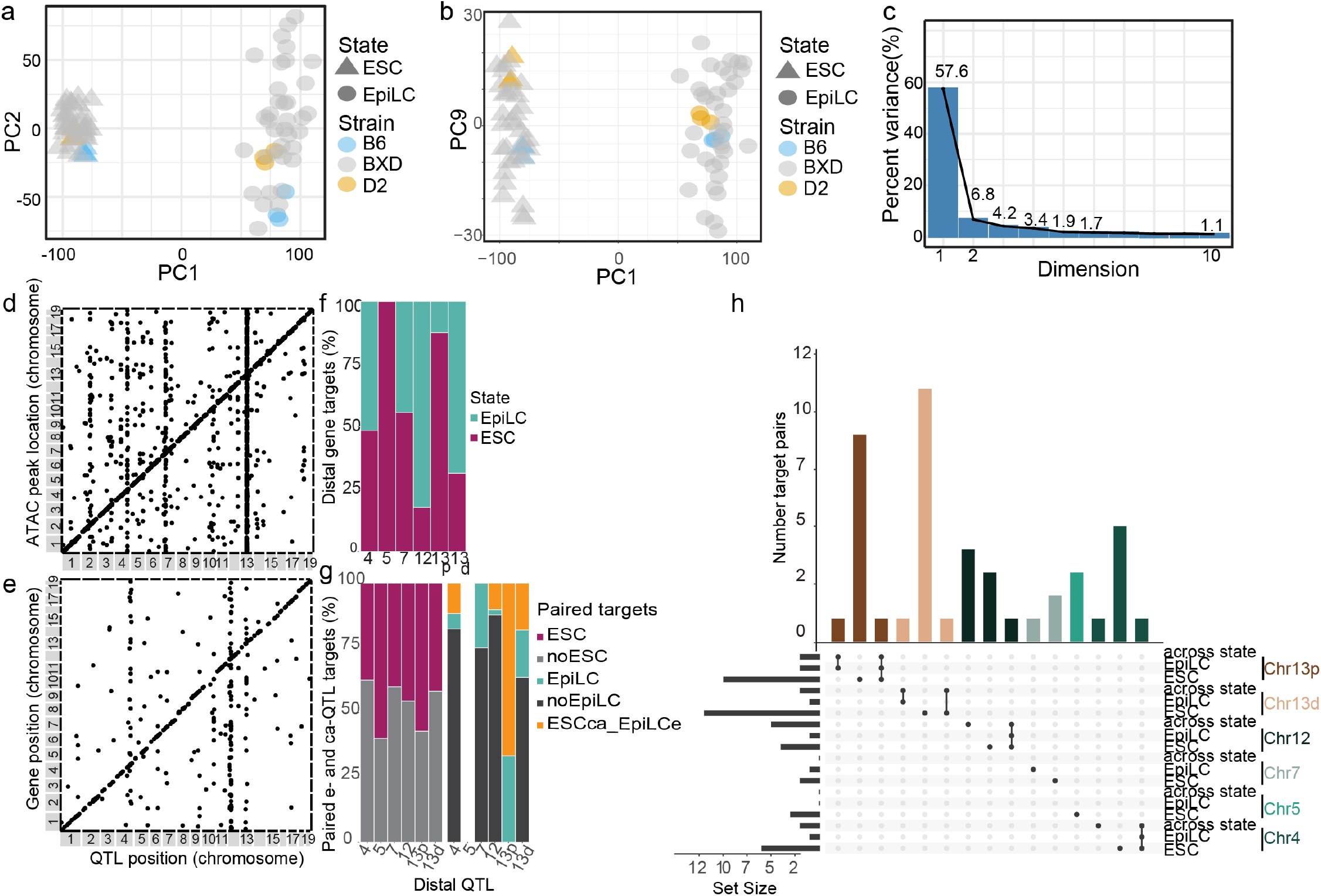
Distal gene targets are paired with proximal chromatin sites co-regulated by hotspot QTL. **a**, Scatterplot of PC1 and PC2 showing variance of total transcript abundance from RNA-seq for 33 distinct strains of BXDs and 3 biological replicates each for B6 and D2 strains in ESCs and EpiLCs. **b**, Similar to *a* showing variance in PC1 versus PC9. **c**, Scree plot shows variance explained for PC1-10. Visual inspection of PC2-9 (total variation = 22.2%) identified that, while biological replicates of parental strains tightly cluster, they are separated by variance captured by the PC, suggesting these PCs represent genetic contribution to transcript abundance within cell state. Individual BXDs can have transcriptomes with greater variation compared to the inbred parents, as is often the case in genetic populations with segregating alleles. **d**, Scatterplot of genomic locations of individual caQTL (x-axis, LOD > 5) versus location of ATAC peak being regulated (y-axis) for EpiLCs. Genetic effects that act locally (n = 4,463), i.e. in *cis*, fall on the diagonal line, while distal acting QTL (n = 1,172) lie off the diagonal. **e**, Similar to *d* plotting eQTL position versus target gene location (n = 318 local-eQTL, n = 200 distal-eQTL). **f**, Proportion of distally regulated genes for each of the 6 *trans-*eQTL between ESCs (maroon) and EpiLCs (teal). The number of target genes regulated by Chrs 4, 7, and 13d distal QTL show roughly equal proportions in both ESC and EpiLC cell states. Chr 5 distal QTL is active exclusively in ESCs, whereas Chrs 12 and 13p distal QTL are biased in regulating more genes in EpiLCs. **g**, Percent of genes targeted by *trans*-eQTL that have matching proximal chromatin accessibility peaks targeted by *trans*-caQTL both that map to the same locus. Matching gene/chromatin pairs were identified by GREAT. Paired targets were determined within state (ESCs= maroon and EpiLCs = teal) and across state (chromatin accessibility in ESCs -> gene expression in EpiLCs = orange). Approximately half of genes controlled by an eQTL in ESCs have a proximally paired chromatin peak regulated by a distal caQTL mapping to the same locus. State-dependent regulation of paired chromatin abundance and gene expression is variable among different QTL. For example, Chr 7 QTL regulates both chromatin accessibility and gene expression at distal targets in the same state; whereas Chr 12 QTL can control chromatin accessibility in ESCs but the expression level of nearby genes maps to the same QTL in EpiLCs. **h**, Upset plot showing relationship between distal targets sharing the same e- and caQTL within or across cell state. Most distal QTL, except Chr 12, show majority of co-regulated targets are present within ESCs (i.e. target chromatin accessibility in ESC is in regulatory neighborhood with target gene, both regulation map back to the same QTL).

**Extended Data Figure 5:**
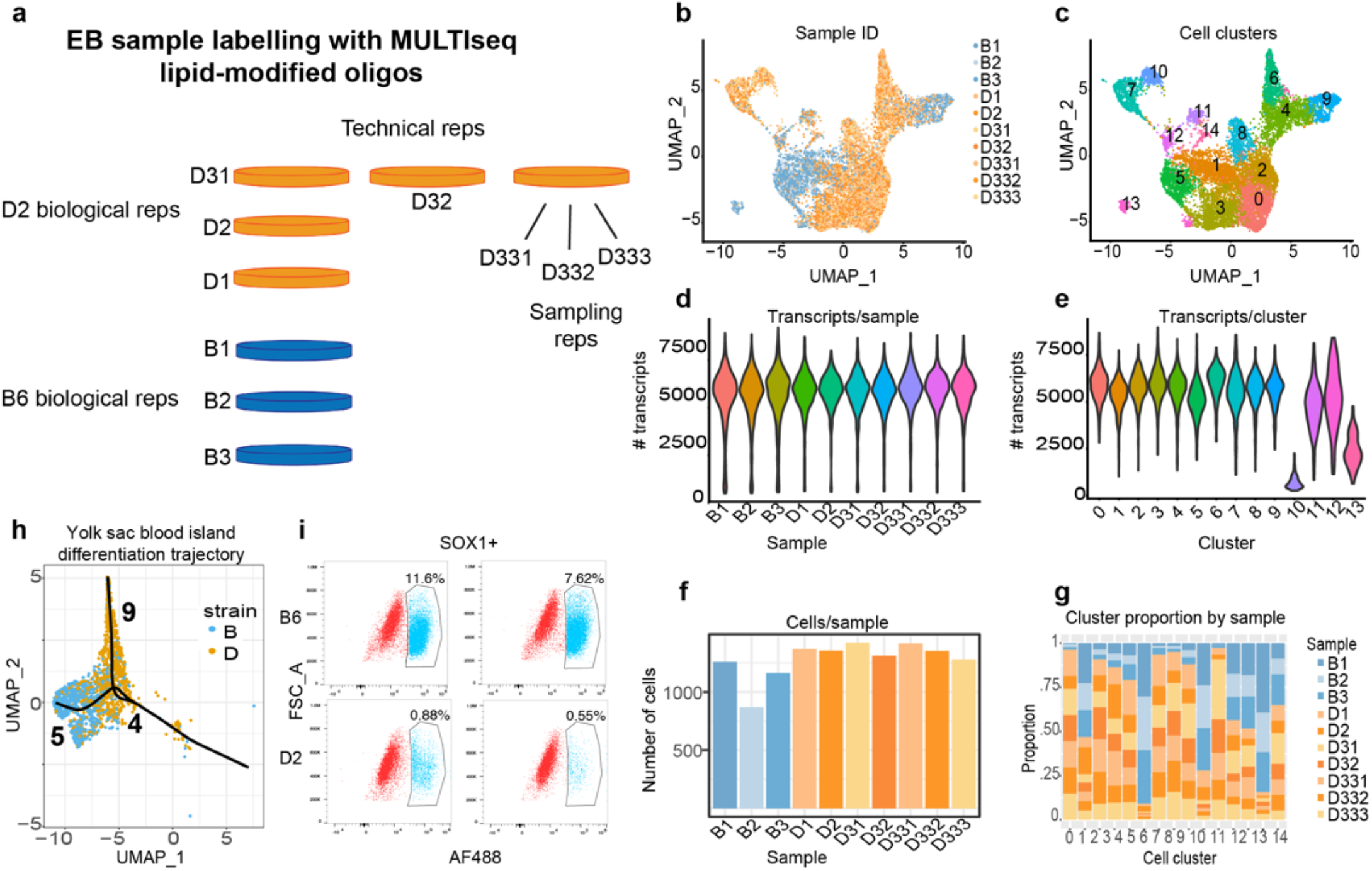
Highly reproducible assay for spontaneous differentiation of ESCs to EBs identifies divergent developmental trajectories between B6 and D2 strains. **a**, Experimental design to assess reproducibility of EB formation and scRNA-seq characterization of cell proportions. Reproducibility was accessed by growing 1) three biological replicates (originating from independently derived ESC lines), 2) one ESC cell line grown for three technical replicates, and 3) sampling replicates (pool of disaggregated EBs sampled three times). Each replicate was barcoded using lipid modified oligonucleotides, pooled, and loaded onto a single 10X Chromium microfluidic lane. **b**, UMAP embeddings for all 10 scRNA-seq libraries colored by individual replicate. Each dot represents a single cell. Cells from replicate EBs clustered in proximity demonstrating reproducibility. **c**, Similar to *b,* UMAP embeddings of cell clustering based on transcript abundance for all 10 samples. **d**, Violin plot of all 10 samples indicating the distribution of number of transcripts per sample. **e,** Violin plot indicating number of transcripts per cluster from 6 samples (3 biological replicates per strain) represented in UMAP shown in Figure 5b. Clusters 10 and 13 had fewer transcripts compared to all other clusters. Attempts at annotation of these two clusters mostly identified cellular processes instead of obvious cellular lineages. **f,** Number of cells per sample shown for each replicate. **g,** Proportion of cells within each cluster color coded by replicate. Overall, biological, technical and sample replicates made up similar proportions of clusters. Because technical and sampling replicates were collected from D2 ESCs, downstream analysis for Figure 5 was performed with only 3 biological replicates for each genotype in order to allow comparisons between similar cell numbers. **h,** UMAP embeddings showing individual cells for clusters 4, 5, and 9 from Figure 5b. Trajectory of pseudotime analysis based on gene expression, using Slingshot, is overlaid on UMAP. Cluster 4 represents yolk sac blood island cells which are common progenitors of vascular endothelium (cluster 5) and primitive erythrocytes (cluster 9). **i**, Scatter plots of replicate FACS analysis on cells from EBs (technical replicate) for both B6 and D2 strains. Cells were labeled with anti-SOX1 antibody. Gated population indicates percentages of SOX1+ cells.

## References

1. Evans, M. J. & Kaufman, M. H. Establishment in culture of pluripotential cells from mouse embryos. Nature 292, 154–156 (1981).

2. Martin, G. R. Isolation of a pluripotent cell line from early mouse embryos cultured in medium conditioned by teratocarcinoma stem cells. Proc Natl Acad Sci USA 78, 7634 (1981).

3. Bradley, A., Evans, M., Kaufman, M. H. & Robertson, E. Formation of germ-line chimaeras from embryo-derived teratocarcinoma cell lines. Nature 309, 255–256 (1984).

4. Buehr, M. & Smith, A. Genesis of embryonic stem cells. Philos Trans R Soc Lond B Biol Sci 358, 1397–1402; discussion 1402 (2003).

5. Nichols, J. et al. Validated germline-competent embryonic stem cell lines from nonobese diabetic mice. Nat Med 15, 814–818 (2009).

6. Czechanski, A. et al. Derivation and characterization of mouse embryonic stem cells from permissive and nonpermissive strains. Nat Protoc 9, 559–574 (2014).

7. Ying, Q.-L. et al. The ground state of embryonic stem cell self-renewal. Nature 453, 519–523 (2008).

8. Osafune, K. et al. Marked differences in differentiation propensity among human embryonic stem cell lines. Nature Biotechnology 26, 313–315 (2008).

9. Li, L., Miu, K.-K., Gu, S., Cheung, H.-H. & Chan, W.-Y. Comparison of multi-lineage differentiation of hiPSCs reveals novel miRNAs that regulate lineage specification. Scientific Reports 8, 9630 (2018).

10. Koyanagi-Aoi, M. et al. Differentiation-defective phenotypes revealed by large-scale analyses of human pluripotent stem cells. Proc Natl Acad Sci USA 110, 20569 (2013).

11. Kyttälä, A. et al. Genetic Variability Overrides the Impact of Parental Cell Type and Determines iPSC Differentiation Potential. Stem Cell Reports 6, 200–212 (2016).

12. Allegrucci, C. & Young, L. E. Differences between human embryonic stem cell lines. Human Reproduction Update 13, 103–120 (2007).

13. Volpato, V. & Webber, C. Addressing variability in iPSC-derived models of human disease: guidelines to promote reproducibility. Dis Model Mech 13, dmm042317 (2020).

14. Skelly, D. A. et al. Mapping the Effects of Genetic Variation on Chromatin State and Gene Expression Reveals Loci That Control Ground State Pluripotency. Cell Stem Cell 27, 459–469.e8 (2020).

15. Ortmann, D. et al. Naive Pluripotent Stem Cells Exhibit Phenotypic Variability that Is Driven by Genetic Variation. Cell Stem Cell 27, 470–481.e6 (2020).

16. Arnold, S. J. & Robertson, E. J. Making a commitment: cell lineage allocation and axis patterning in the early mouse embryo. Nat Rev Mol Cell Biol 10, 91–103 (2009).

17. Hackett, J. A. & Surani, M. A. Regulatory Principles of Pluripotency: From the Ground State Up. Cell Stem Cell 15, 416–430 (2014).

18. Nichols, J. & Smith, A. Naive and primed pluripotent states. Cell Stem Cell 4, 487–492 (2009).

19. Gardner, R. L. & Beddington, R. S. Multi-lineage ‘stem’ cells in the mammalian embryo. J Cell Sci Suppl 10, 11–27 (1988).

20. Brons, I. G. M. et al. Derivation of pluripotent epiblast stem cells from mammalian embryos. Nature 448, 191–195 (2007).

21. Hayashi, K., Ohta, H., Kurimoto, K., Aramaki, S. & Saitou, M. Reconstitution of the Mouse Germ Cell Specification Pathway in Culture by Pluripotent Stem Cells. Cell 146, 519–532 (2011).

22. Morgani, S., Nichols, J. & Hadjantonakis, A.-K. The many faces of Pluripotency: in vitro adaptations of a continuum of in vivo states. BMC Developmental Biology 17, 7 (2017).

23. Smith, A. Formative pluripotency: the executive phase in a developmental continuum. Development 144, 365 (2017).

24. Kawase, E. et al. Strain difference in establishment of mouse embryonic stem (ES) cell lines. Int J Dev Biol 38, 385–390 (1994).

25. Sharova, L. V. et al. Global gene expression profiling reveals similarities and differences among mouse pluripotent stem cells of different origins and strains. Dev Biol 307, 446–459 (2007).

26. Garbutt, T. A. et al. Permissiveness to form pluripotent stem cells may be an evolutionarily derived characteristic in Mus musculus. Scientific Reports 8, 14706 (2018).

27. Schnabel, L. V., Abratte, C. M., Schimenti, J. C., Southard, T. L. & Fortier, L. A. Genetic background affects induced pluripotent stem cell generation. Stem Cell Research & Therapy 3, 30 (2012).

28. Hackett, J. A., Kobayashi, T., Dietmann, S. & Surani, M. A. Activation of Lineage Regulators and Transposable Elements across a Pluripotent Spectrum. Stem Cell Reports 8, 1645–1658 (2017).

29. Bernstein, B. E. et al. A bivalent chromatin structure marks key developmental genes in embryonic stem cells. Cell 125, 315–326 (2006).

30. Bonifer, C. & Cockerill, P. N. Chromatin priming of genes in development: Concepts, mechanisms and consequences. Experimental Hematology 49, 1–8 (2017).

31. Pękowska, A. et al. Gain of CTCF-Anchored Chromatin Loops Marks the Exit from Naive Pluripotency. Cell Syst 7, 482–495.e10 (2018).

32. Factor, D. C. et al. Epigenomic comparison reveals activation of ‘seed’ enhancers during transition from naive to primed pluripotency. Cell Stem Cell 14, 854–863 (2014).

33. Novo, C. L. et al. Long-Range Enhancer Interactions Are Prevalent in Mouse Embryonic Stem Cells and Are Reorganized upon Pluripotent State Transition. Cell Rep 22, 2615–2627 (2018).

34. Chen, T. & Dent, S. Y. R. Chromatin modifiers and remodellers: regulators of cellular differentiation. Nat Rev Genet 15, 93–106 (2014).

35. Yadav, T., Quivy, J.-P. & Almouzni, G. Chromatin plasticity: A versatile landscape that underlies cell fate and identity. Science 361, 1332 (2018).

36. Catarino, R. R. & Stark, A. Assessing sufficiency and necessity of enhancer activities for gene expression and the mechanisms of transcription activation. Genes Dev 32, 202–223 (2018).

37. Buecker, C. et al. Reorganization of enhancer patterns in transition from naive to primed pluripotency. Cell Stem Cell 14, 838–853 (2014).

38. Yang, S.-H. et al. ZIC3 Controls the Transition from Naive to Primed Pluripotency. Cell Reports 27, 3215–3227.e6 (2019).

39. Ito, K. & Suda, T. Metabolic requirements for the maintenance of self-renewing stem cells. Nat Rev Mol Cell Biol 15, 243–256 (2014).

40. Van Winkle, L. J. & Ryznar, R. One-Carbon Metabolism Regulates Embryonic Stem Cell Fate Through Epigenetic DNA and Histone Modifications: Implications for Transgenerational Metabolic Disorders in Adults. Frontiers in Cell and Developmental Biology 7, 300 (2019).

41. Gonzales, K. A. U. et al. Deterministic Restriction on Pluripotent State Dissolution by Cell-Cycle Pathways. Cell 162, 564–579 (2015).

42. ten Berge, D. et al. Embryonic stem cells require Wnt proteins to prevent differentiation to epiblast stem cells. Nat Cell Biol 13, 1070–1075 (2011).

43. de Jaime-Soguero, A., Abreu de Oliveira, W. A. & Lluis, F. The Pleiotropic Effects of the Canonical Wnt Pathway in Early Development and Pluripotency. Genes (Basel) 9, 93 (2018).

44. Jiang, Y. et al. Zinc finger E-box-binding homeobox 1 (ZEB1) is required for neural differentiation of human embryonic stem cells. J Biol Chem 293, 19317–19329 (2018).

45. Wang, Q. et al. The p53 Family Coordinates Wnt and Nodal Inputs in Mesendodermal Differentiation of Embryonic Stem Cells. Cell Stem Cell 20, 70–86 (2017).

46. Venere, M. et al. Sox1 marks an activated neural stem/progenitor cell in the hippocampus. Development 139, 3938 (2012).

47. Peirce, J. L., Lu, L., Gu, J., Silver, L. M. & Williams, R. W. A new set of BXD recombinant inbred lines from advanced intercross populations in mice. BMC Genetics 5, 7 (2004).

48. Friedman, J. R. et al. KAP-1, a novel corepressor for the highly conserved KRAB repression domain. Genes Dev 10, 2067–2078 (1996).

49. Baker, C. L. et al. Tissue-Specific <em>Trans</em> Regulation of the Mouse Epigenome. Genetics 211, 831 (2019).

50. Palis, J., Robertson, S., Kennedy, M., Wall, C. & Keller, G. Development of erythroid and myeloid progenitors in the yolk sac and embryo proper of the mouse. Development 126, 5073–5084 (1999).

51. Bedzhov, I. & Zernicka-Goetz, M. Self-organizing properties of mouse pluripotent cells initiate morphogenesis upon implantation. Cell 156, 1032–1044 (2014).

52. Neagu, A. et al. In vitro capture and characterization of embryonic rosette-stage pluripotency between naive and primed states. Nat Cell Biol 22, 534–545 (2020).

53. Argelaguet, R. et al. Multi-omics profiling of mouse gastrulation at single-cell resolution. Nature 576, 487–491 (2019).

54. Maurano, M. T. et al. Systematic Localization of Common Disease-Associated Variation in Regulatory DNA. Science 337, 1190 (2012).

55. Johnson, K. R., Lane, P. W., Ward-Bailey, P. & Davisson, M. T. Mapping the mouse dactylaplasia mutation, Dac, and a gene that controls its expression, mdac. Genomics 29, 457–464 (1995).

56. Plamondon, J. A., Harris, M. J., Mager, D. L., Gagnier, L. & Juriloff, D. M. The clf2 gene has an epigenetic role in the multifactorial etiology of cleft lip and palate in the A/WySn mouse strain. Birth Defects Res A Clin Mol Teratol 91, 716–727 (2011).

57. Treger, R. S. et al. The Lupus Susceptibility Locus Sgp3 Encodes the Suppressor of Endogenous Retrovirus Expression SNERV. Immunity 50, 334–347.e9 (2019).

58. Bruno, M., Mahgoub, M. & Macfarlan, T. S. The Arms Race Between KRAB–Zinc Finger Proteins and Endogenous Retroelements and Its Impact on Mammals. Annu. Rev. Genet. 53, 393–416 (2019).

59. Kauzlaric, A. et al. The mouse genome displays highly dynamic populations of KRAB-zinc finger protein genes and related genetic units. PLoS One 12, e0173746–e0173746 (2017).

60. Elmer, J. L. & Ferguson-Smith, A. C. Strain-Specific Epigenetic Regulation of Endogenous Retroviruses: The Role of Trans-Acting Modifiers. Viruses 12, 810 (2020).

61. Smukler, S. R., Runciman, S. B., Xu, S. & van der Kooy, D. Embryonic stem cells assume a primitive neural stem cell fate in the absence of extrinsic influences. J Cell Biol 172, 79–90 (2006).

62. Andoniadou, C. L. & Martinez-Barbera, J. P. Developmental mechanisms directing early anterior forebrain specification in vertebrates. Cell Mol Life Sci 70, 3739–3752 (2013).

63. Lenka, N. & Ramasamy, S. K. Neural induction from ES cells portrays default commitment but instructive maturation. PLoS One 2, e1349–e1349 (2007).

64. The GTEx Consortium atlas of genetic regulatory effects across human tissues. Science 369, 1318 (2020).

65. Corces, M. R. et al. An improved ATAC-seq protocol reduces background and enables interrogation of frozen tissues. Nature Methods 14, 959–962 (2017).

66. Corces, M. R. et al. Lineage-specific and single-cell chromatin accessibility charts human hematopoiesis and leukemia evolution. Nat Genet 48, 1193–1203 (2016).

67. Baker, C. L. et al. PRDM9 drives evolutionary erosion of hotspots in Mus musculus through haplotype-specific initiation of meiotic recombination. PLoS Genet 11, e1004916 (2015).

68. McGinnis, C. S. et al. MULTI-seq: sample multiplexing for single-cell RNA sequencing using lipid-tagged indices. Nat Methods 16, 619–626 (2019).

69. Langmead, B., Trapnell, C., Pop, M. & Salzberg, S. L. Ultrafast and memory-efficient alignment of short DNA sequences to the human genome. Genome Biology 10, R25 (2009).

70. Raghupathy, N. et al. Hierarchical analysis of RNA-seq reads improves the accuracy of allele-specific expression. Bioinformatics 34, 2177–2184 (2018).

71. Robinson, M. D., McCarthy, D. J. & Smyth, G. K. edgeR: a Bioconductor package for differential expression analysis of digital gene expression data. Bioinformatics 26, 139–140 (2010).

72. Yalcin, B. et al. Sequence-based characterization of structural variation in the mouse genome. Nature 477, 326–329 (2011).

73. Keane, T. M. et al. Mouse genomic variation and its effect on phenotypes and gene regulation. Nature 477, 289–294 (2011).

74. Pertea, M., Kim, D., Pertea, G. M., Leek, J. T. & Salzberg, S. L. Transcript-level expression analysis of RNA-seq experiments with HISAT, StringTie and Ballgown. Nature Protocols 11, 1650–1667 (2016).

75. Zhang, Y. et al. Model-based Analysis of ChIP-Seq (MACS). Genome Biology 9, R137 (2008).

76. Quinlan, A. R. & Hall, I. M. BEDTools: a flexible suite of utilities for comparing genomic features. Bioinformatics 26, 841–842 (2010).

77. Broman, K. W. et al. R/qtl2: Software for Mapping Quantitative Trait Loci with High-Dimensional Data and Multiparent Populations. Genetics 211, 495–502 (2019).

78. Yu, G., Wang, L.-G. & He, Q.-Y. ChIPseeker: an R/Bioconductor package for ChIP peak annotation, comparison and visualization. Bioinformatics 31, 2382–2383 (2015).

79. McLean, C. Y. et al. GREAT improves functional interpretation of cis-regulatory regions. Nature Biotechnology 28, 495–501 (2010).

80. Subramanian, A. et al. Gene set enrichment analysis: A knowledge-based approach for interpreting genome-wide expression profiles. Proc Natl Acad Sci USA 102, 15545 (2005).

81. Mootha, V. K. et al. PGC-1α-responsive genes involved in oxidative phosphorylation are coordinately downregulated in human diabetes. Nature Genetics 34, 267–273 (2003).

82. Liberzon, A. et al. Molecular signatures database (MSigDB) 3.0. Bioinformatics 27, 1739–1740 (2011).

83. Liberzon, A. et al. The Molecular Signatures Database (MSigDB) hallmark gene set collection. Cell Syst 1, 417–425 (2015).

84. Leng, N. et al. EBSeq-HMM: a Bayesian approach for identifying gene-expression changes in ordered RNA-seq experiments. Bioinformatics 31, 2614–2622 (2015).

85. Gu, Z., Eils, R. & Schlesner, M. Complex heatmaps reveal patterns and correlations in multidimensional genomic data. Bioinformatics 32, 2847–2849 (2016).

86. Yu, G., Wang, L.-G., Han, Y. & He, Q.-Y. clusterProfiler: an R Package for Comparing Biological Themes Among Gene Clusters. OMICS: A Journal of Integrative Biology 16, 284–287 (2012).

87. Schep, A. N., Wu, B., Buenrostro, J. D. & Greenleaf, W. J. chromVAR: inferring transcription-factor-associated accessibility from single-cell epigenomic data. Nature Methods 14, 975–978 (2017).

88. Mei, S. et al. Cistrome Data Browser: a data portal for ChIP-Seq and chromatin accessibility data in human and mouse. Nucleic Acids Research 45, D658–D662 (2017).

89. Zheng, R. et al. Cistrome Data Browser: expanded datasets and new tools for gene regulatory analysis. Nucleic Acids Research 47, D729–D735 (2019).

90. Sheffield, N. C. & Bock, C. LOLA: enrichment analysis for genomic region sets and regulatory elements in R and Bioconductor. Bioinformatics 32, 587–589 (2016).

91. Stuart, T. et al. Comprehensive Integration of Single-Cell Data. Cell 177, 1888–1902.e21 (2019).

92. Nowotschin, S. et al. The emergent landscape of the mouse gut endoderm at single-cell resolution. Nature 569, 361–367 (2019).

93. Motenko, H., Neuhauser, S. B., O’Keefe, M. & Richardson, J. E. MouseMine: a new data warehouse for MGI. Mamm Genome 26, 325–330 (2015).

94. Street, K. et al. Slingshot: cell lineage and pseudotime inference for single-cell transcriptomics. BMC Genomics 19, 477 (2018).

